# *nubbin*, *ventral veinless*, and *pdm3* play diverse roles in butterfly wing pattern development

**DOI:** 10.64898/2026.08.06.742654

**Authors:** Jeanne M. C. McDonald, Qin Guo, Seydeanna Delgado, Connor A. Amendola, Iya A. Garg, Robert D. Reed

## Abstract

Butterfly wings present a tremendous gallery of colorful patterns, offering a unique opportunity to study how developmental pattern formation processes evolve. We still do not understand the genetic basis of several key aspects of wing pattern development, however. Three paralogous POU domain transcription factors *nubbin, ventral veinless* (*vvl*), and *pdm3* are all known wing development genes in *Drosophila melanogaster*. Here we combine gene expression and knockout approaches to show that each of these genes plays multiple novel wing patterning roles in the common buckeye butterfly, *Junonia coenia*. We found that *nubbin* controls eyespot pattern determination via a non-cell autonomous repressor-like effect originating at the wing veins, such that *nubbin* knockouts have larger eyespots. *nubbin* also regulates pigment identity and scale morphology across the wing. We also found that *vvl* regulates pigment identity of the discal bands and ventral hindwing. Last, we found that *pdm3* is required for determining the outer rings of eyespot patterns, where it is co-expressed with *spalt* and the lncRNA *ivory. pdm3* is also necessary for determining wing margin stripes, where it is again co-expressed with *spalt*, leading us to propose that the eyespot and wing margin gene regulatory networks could be homologous. Together, our work shows that POU domain transcription factors play diverse roles in butterfly wing pattern development and highlights *nubbin* as one of the first genes implicated in the repressive function of wing veins in color pattern determination.

**Highlights:**

- *nubbin* regulates eyespot development, likely via a repressor from the wing veins
- *nubbin* controls scale color and morphology across the wing
- *vvl* regulates color pattern readout and vein development
- *pdm3* coordinates eyespot development and is co-expressed with *spalt* and *ivory*
- Expression of genes in the eyespot and wing margin suggests network homology

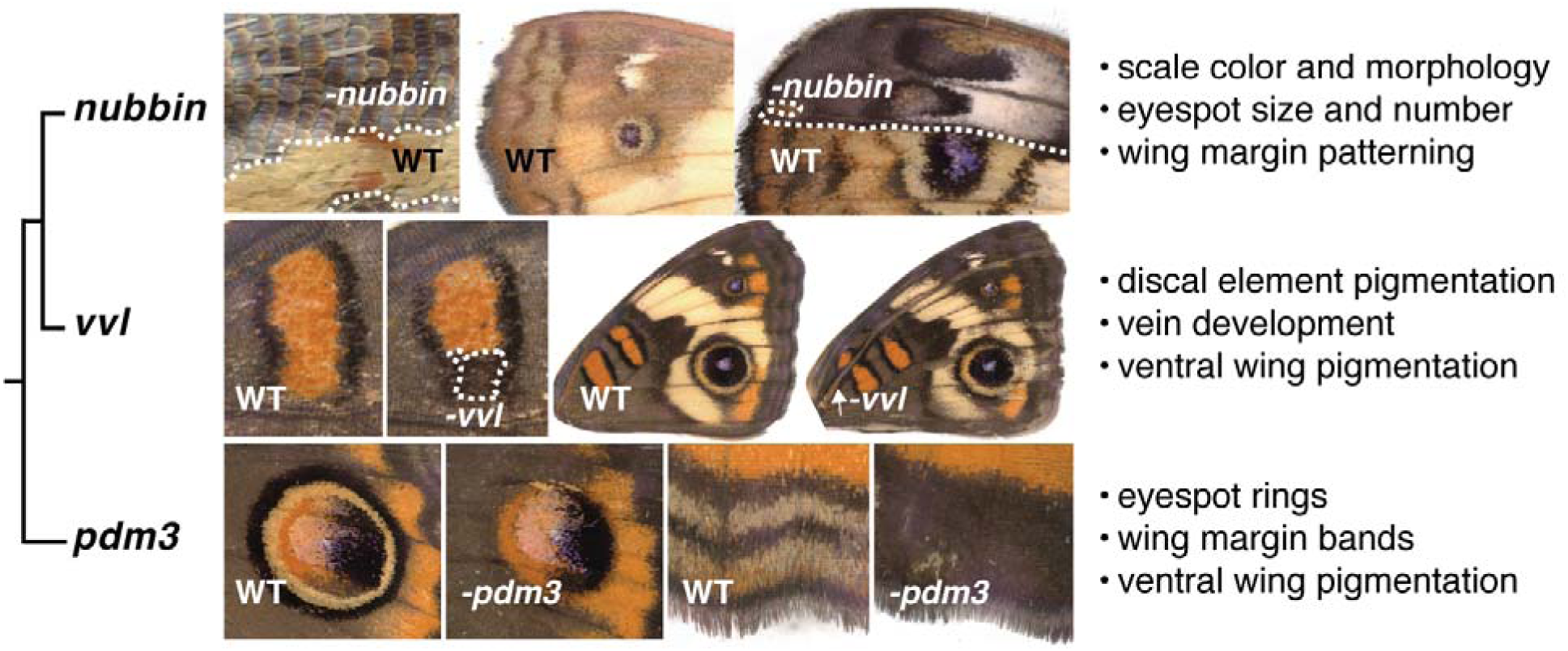

## 1. Introduction

The rich diversity and experimental tractability of butterfly wing patterns makes them an ideal system for studies in evolutionary developmental biology. Specifically, the butterfly family Nymphalidae displays wing patterns that are variations upon a set of homologous pattern elements (Fig. 1A-B) that allow us to ask what kind of developmental genetic changes underlie pattern evolution at different phylogenetic scales. A growing number of genes have been functionally linked to some of these pattern elements across a few species (see Fig. 1C for an example), but we still lack genes for some of the key elements and processes. To reconstruct how color patterns evolved across species, we need to generate a more complete developmental model of the nymphalid groundplan, including characterizing additional patterning genes.

**Fig. 1:**
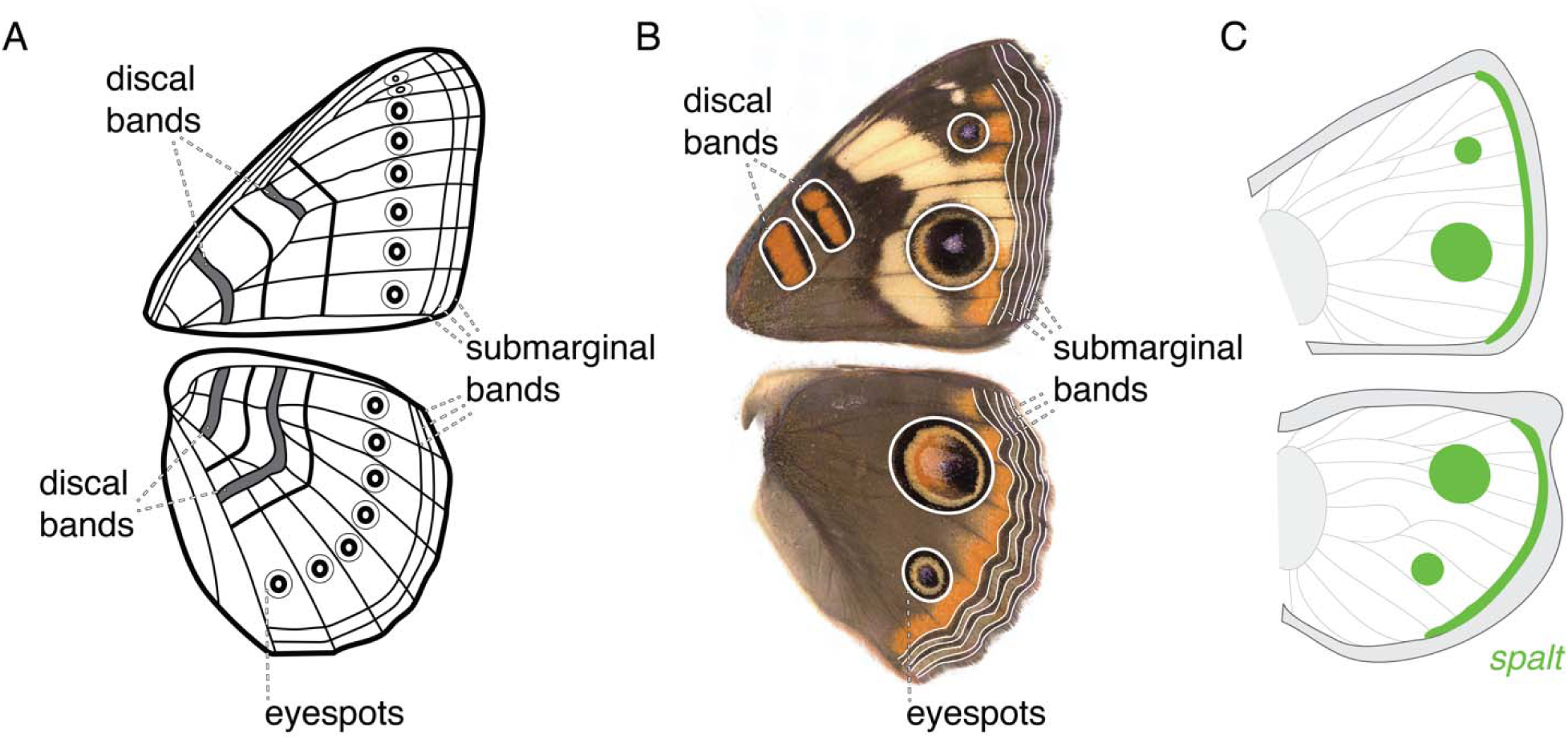
Key wing pattern elements of the nymphalid groundplan and associated gene expression in *J. coenia.* The nymphalid groundplan, as adapted from (Schwanwitsch, 1929), represents a set of pattern elements that can be homologized across nymphalid species. Key pattern elements for this study are highlighted on (A) the groundplan and in (B) *J. coenia*. (C) An example of a known wing patterning gene, *spalt,* that is expressed in the eyespots and wing margins in developing pupal wing imaginal disks. *spalt’s* function in patterning these traits was previously validated using CRISPR (Zhang and Reed, 2016).

In this study, we investigated three paralogous POU domain genes that are known to regulate wing development in other species: *nubbin*, *ventral veinless* (*vvl*), and *pou domain motif 3* (*pdm3*). We assessed the functions of these genes in a model nymphalid butterfly, the buckeye butterfly *Junonia coenia,* in part to set the stage for future comparative studies across Nymphalidae. *nubbin* controls wing pattern and growth in many insects, including flies (Ng, Diaz-Benjumea, and Cohen 1995), crickets (Lim, 2009), and beetles (Tomoyasu et al., 2009), and its expression is associated with wing scale development in the orange sulphur butterfly *Colias eurytheme* (Loh et al., 2025b) and with abdominal pigmentation regulation in bumblebees (Hines et al., 2025). *vvl* is necessary for wing vein differentiation and wing growth in *Drosophila melanogaster* (Certel et al., 2000; J. F. de Celis et al., 1995). Differences in the distal black forewing pattern in the red postman butterfly, *Heliconius erato,* are also associated with divergence in a genomic region that includes the *vvl* gene (Van Belleghem et al., 2017). A third POU domain factor associated with fly wing development is *pdm3*, which is expressed in the wing notum and hinge during *D. melanogaster* larval development (Everetts et al., 2021). *pdm3* also suppresses melanin pigmentation in *D. melanogaster* abdomens and has been linked to repeated evolution in abdominal pigmentation in a *Drosophila* clade (Rogers et al., 2014; Yassin et al., 2016). Excitingly, *pdm3* was also recently discovered to regulate wing patterning in the painted lady butterfly, *Vanessa cardui* (Loh et al., 2025a).

Here, we investigated whether *nubbin, vvl,* and *pdm3* have evolved to regulate wing color patterning in the common buckeye butterfly, *Junonia coenia,* using in situ hybridization and CRISPR-Cas9 mutation. We found each of these genes plays multiple roles in wing color pattern development: *nubbin* controls both scale color and morphology across the wing, *vvl* coordinates the pigmentation readout for multiple wing pattern elements, and *pdm3* is required for patterning a specific subset of eyespot and wing margin pattern features. This work not only identifies new genes involved in butterfly wing patterning, but sheds new light on the process of eyespot development, which has been a particular focus of study in butterflies as a model novel trait (Beldade and Monteiro, 2021). We provide the first evidence to suggest a signal originating from the wing veins downstream of *nubbin* non-cell autonomously regulates eyespot development. We also show *pdm3* is co-expressed with known eyespot patterning genes *spalt* and *ivory* and suggest that it is downstream of the eyespot morphogen based on the temporal progression of its expression.

## 2. Material and Methods

### 2.1 Butterfly rearing

The *J. coenia* colony was reared on an artificial diet (Nijhout, 1980; Yamamoto, 1969) at 27°C, 70% humidity, and a 16hr : 8hr light : dark cycle.

### 2.2 Phylogenetic reconstruction

We constructed two gene phylogenies for this clade of paralogous POU domain factors: one to identify orthologs for *nubbin, vvl,* and *pdm3* across holometabolous insects (*Junonia coenia, Vanessa atalanta*, *Bombyx mori, Aedes aegypti, Tribolium castaneum, Cheumatopsyche charites,* and *Drosophila melanogaster)* to confirm our gene annotations in *J. coenia,* and one to confirm these orthology assignments by including additional species across Pancrustacea and an outgroup mollusc species (*Aphis gossypii, Folsomia candida, Daphnia pulicaria, Caligus rogercresseyi,* and *Crassostrea virginica*).

We identified the top hit for each *D. melanogaster* protein sequence for nubbin, vvl, and pdm3 on NCBI BLAST for the following species: *Vanessa atalanta*, *Bombyx mori, Aedes aegypti, Tribolium castaneum, Aphis gossypii, Folsomia candida, Daphnia pulicaria, Caligus rogercresseyi,* and *Crassostrea virginica*. For *J. coenia,* we first performed a BLAST search using the Junonia_coenia_v2 reference genome from Lepbase (van der Burg et al., 2020). The *J. coenia* sequences were manually re-annotated due to fragmentation in the original annotations using Iso-seq data from (Fandino et al., 2024). The gene annotation for *nubbin* and *pdm3* are reported using accession numbers from Junonia_coenia_v2 (van der Burg et al., 2020) and the gene annotation for *vvl* is reported using the accession number from the new annotation in (Fandino et al., 2024). For *Cheumatopsyche charites,* we performed a BLAST search using the reference genome from (Ge et al., 2022). We also selected the *D. melanogaster* sequence for acj6, a POU domain factor that is not expressed in the butterfly wing, to root the tree.

We then used MAFFT with default settings for amino acid sequences to generate a multiple sequence alignment (Katoh and Standley, 2013). We assessed this alignment in Jalview (Waterhouse et al., 2009). Then, using IQ-TREE 2 (Minh et al., 2020), we determined the best substitution model was Q.insect+F+I+G4 using ModelFinder and then generated a maximum likelihood tree using 1000 bootstrap replicates.

All code and protein sequences are available at: https://github.com/jeannexpression/pdm_tree

### 2.3 RNA-seq analysis

We re-aligned RNA-seq data collected from wings at four different developmental stages: fifth instar, prepupa, day 3 pupae, and day 7 pupae from (van der Burg et al., 2019) to jcgen_v2 with our updated annotation file using STAR v.2.7.11b (Dobin et al., 2013) using default parameters except for --alignIntronMin 5, --alignIntronMax 50000, and --genomeSAindexNbases 13. Transcripts were quantified using the featureCounts program from the Subread package (Liao et al., 2014). We used the plotCounts function from the package DESeq2 (Love et al., 2014) and ggplot2 (Wickham, 2016) in R version 4.4.3 to plot normalized counts for each gene across developmental stages.

### 2.4 In situ hybridization

We performed in situ hybridization using hybridization chain reaction (HCR) v3.0 to visualize expression of *nubbin, vvl,* and *pdm3* following published protocols (Bruce et al., 2021; Choi et al., 2018). Wing disks from fifth instar larvae and hindwing disks from day 1, day 2, and day 3 pupae were dissected, fixed at room temperature in 9.25% formaldehyde (Thermo Fisher Scientific Pierce 16% w/v), and stored in methanol for up to two months beforehand. Day 1 pupal wings were dissected between 7-18h post-pupation (pupal development takes an average of 178h, so this corresponds to 4-10% pupal development); day 2 wings were dissected 31-40h post-pupation (17-22%), and day 3 wings were dissected at 52-63h post-pupation (29-35%). All probes were ordered from Molecular Instruments. *J. coenia nubbin, vvl,* and *pdm3* probes were in the B1 channel. *spalt* probes were in the B3 channel. *ivory* probes were in the B2 channel. We used B1 647nm hairpins and B3 546nm hairpins for in situs for *pdm3+spalt, nubbin+spalt,* and *vvl+spalt* (Molecular Instruments). We used B1 647nm hairpins and B2 546nm hairpins for *pdm3+ivory* in situs. DAPI nuclear marker (AAT Bioquest) was used as a counterstain for all in situs. Negative controls were performed using wing disks at the same developmental stages without probes but with the same hairpins. Wing disks were mounted in ProLong Gold antifade reagent (Thermo Fisher Scientific) and imaged within two weeks. We used a Zeiss LSM880 confocal/multiphoton microscope with Zen software to image the wings. We used the tile scan feature to image larger fifth instar and pupal wing disks. Negative control disks were imaged re-using the same imaging settings. .czi files were then processed in ImageJ.

### 2.5 CRISPR-Cas9

We followed a previously published protocol to generate mosaic knockouts (mKOs) using CRISPR-Cas9 for each of the three genes, *nubbin*, *vvl,* and *pdm3* in *J. coenia* (Zhang and Reed, 2017). All eggs were injected between 1-4 hours after egg lay. Each gene was targeted with two sgRNAs that were designed to exon sequences (Table 1) using Geneious software. We selected sgRNAs with zero predicted off-target sites and with an activity score greater than 0.4 according to (Doench et al., 2016). sgRNAs were then synthesized by Integrated DNA Technologies and injected at 150-272 ng/ul. Alt-R S.p. Hi-Fi Cas9 Nuclease V3 (Integrated DNA Technologies) was injected at a concentration of 500 ng/ul. sgRNAs and Cas9 were diluted with nuclease-free water. sgRNA-Cas9 solutions prepared for injections were dyed with a small amount of amaranth (Sigma-Aldrich) to aid with injections.

**Table 1:**
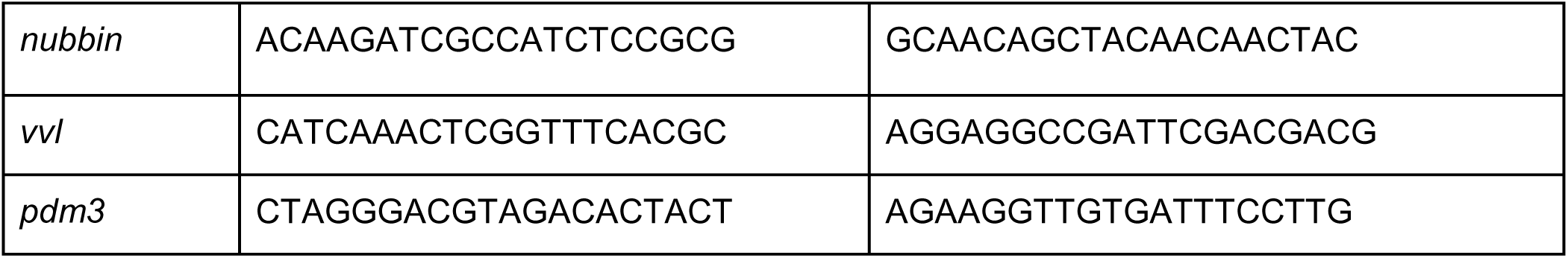
gRNA sequences.

For *nubbin*, 1363 eggs were injected with two gRNAs at a concentration between 150-250 ng/ul, with an average hatch rate of 17%. About 61% of larvae survived to adulthood. Of these, we found a mutant phenotype in 15 butterflies, or 9% of adults.

For *vvl,* 793 eggs were injected with two gRNAs at a concentration of 250-272 ng/ul, with an average hatch rate of 35%. On average, 34% of larvae survived to adulthood. 12 butterflies had a mutant wing phenotype, or 16% of all adults that eclosed.

For *pdm3*, 440 eggs were injected with two gRNAs at a concentration of 200-250 ng/ul, with an average hatch rate of 37%. 40% of larvae survived to adulthood. 60 butterflies had a mutant wing phenotype, or 89% of all adults that eclosed. There were no sex-specific differences in survivorship and mutant phenotype, as has been observed in *pdm3* knockouts in *V. cardui* (Loh et al., 2025a).

Differences in hatch rate, survivorship, and mutation rate between *nubbin, vvl,* and *pdm3* gene knockouts could be due to differences in injection technique and larval rearing rather than differences in lethality of the gene knockouts.

Wings from mKOs were dissected and then imaged using a Keyence VHX-7000. Additional mKO images are available in Supp. File 5 and mKO phenotypes are summarized in Table S1.

### 2.6 Genotyping CRISPR mutants

We genotyped selected mKO butterflies (nubbin_02, nubbin_04, nubbin_67, pdm3_12, pdm3_45, vvl_20) to confirm mutation of each target gene. Sequencing primers were designed at least 130 bp away from the CRISPR-Cas9 cut sites using Geneious (Table 2). We confirmed primer specificity using BLAST. DNA extraction and PCR amplification of target regions were performed using Phire Tissue Direct PCR Master Mix (Thermo Fisher Scientific). We confirmed PCR amplification using gel electrophoresis. We purified PCR products using ExoSAP-IT (Thermo Fisher Scientific). Sanger sequencing was performed by Cornell University’s Genomics Facility (*nubbin, pdm3* mKOs) and by Eurofins Scientific (*vvl* mKOs). Results were analyzed using Synthego’s Inference of CRISPR Edits tool (EditCo Bio, 2026) and confirmed successful mutation for each target (Supp. File 3).

**Table 2:**
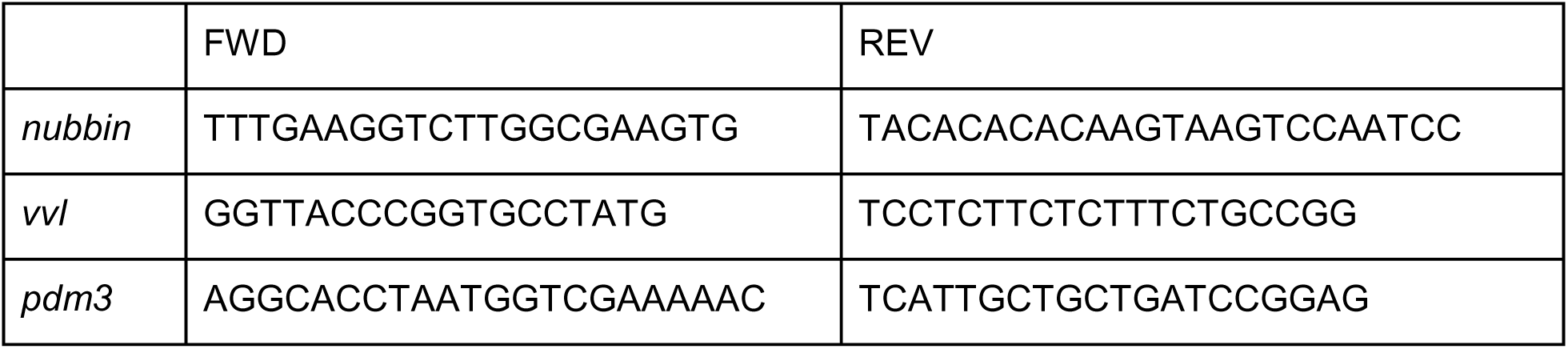
Sequencing primers.

## 3. Results

### 3.1 Phylogenetic and gene expression confirmation of nubbin, vvl, and pdm3 orthologs in J. coenia wings

*nubbin*, *vvl*, and *pdm3* are three genes described in *D. melanogaster* that each belong to ancient families of POU domain factors. The POU3 class (*vvl*) and POU6 class (*pdm3*) are thought to have evolved before the last common ancestor of all animals, and the POU2 class (*nubbin*) evolved in bilaterians (Gold et al., 2014). To confirm our gene annotations for *nubbin*, *vvl*, and *pdm3* in *J. coenia,* we constructed a maximum likelihood phylogeny with 1000 bootstraps using the top BLAST hits for nubbin, vvl, and pdm3 protein sequences from *D. melanogaster* to identify orthologs for each of these proteins in *J. coenia* and other insects (Supp. File 1, Fig. S1). This analysis confirmed our annotations. We also constructed a phylogeny across Pancrustacea, which confirmed these results (Supp. File 1, Fig. S2). We then used previously published RNA sequencing data to confirm that these three genes are expressed in the developing wing disks in *J. coenia* (van der Burg et al., 2019). These data show all three genes are expressed during key timepoints for color pattern development in the wing (Supp. File 2).

### 3.2 nubbin regulates wing color pattern formation, pigmentation, and scale morphology

We characterized the expression of *nubbin* in *J. coenia* using in situ hybridization to determine whether this gene is expressed in any spatially or temporally restricted patterns, using *spalt* expression as a marker for eyespot and wing margin color pattern elements. We found *nubbin* expression correlated with the development of the wing veins, with expression detected along the lacunae, the gaps between the dorsal and ventral epithelial layers where the wing veins form, throughout fifth instar. We continued to observe *nubbin* expression in the developing veins during pupal development (Fig. 2). *nubbin* was also expressed in a proximal band of the wing that intersects with the origin points of the lacunae in fifth instar and pupal wing disks. In day 3 pupae we observed that *nubbin* was also expressed ubiquitously across the wing, although at a lower expression level than in the wing veins (Fig. 2). These data suggest that *nubbin* could play a role in wing vein specification and potentially wing color pattern formation.

**Fig. 2:**
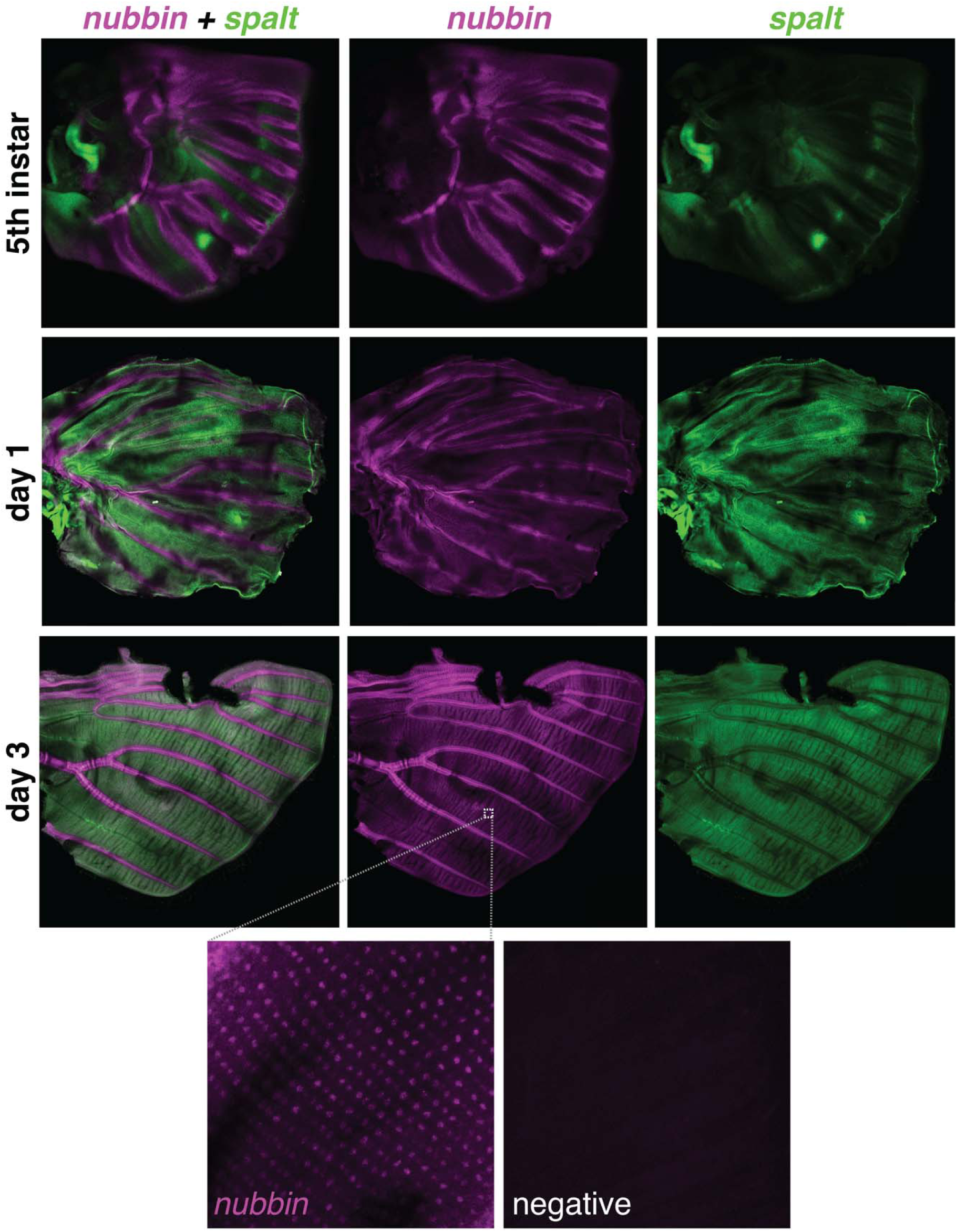
nubbin is upregulated in the developing wing veins and ubiquitously expressed in the pupal wing. During 5th instar, *nubbin* is expressed along the developing wing veins, i.e. the lacunae. In day 1 and day 3 pupae, *nubbin* is still expressed at the lacunae. In day 3 pupae, both *nubbin* and *spalt* are expressed throughout the wing disk. *nubbin* is also expressed in the basal band of the wing that intersects the origins of the lacunae.

We then determined whether *nubbin* is necessary for wing patterning by knocking it out using CRISPR-Cas9 to produce deletion mosaics. These knockouts dramatically transformed wing color patterns, showing effects on pigmentation, scale structure, and color pattern formation (Fig. 3, Supp. File 5). We found *nubbin* mKOs eliminated orange ommochrome pigments broadly across the wings, resulting in depigmented scales (Fig. 3B-C, E, G-H), as well as scales with darker black and gray melanins (Fig. 3B-C, E, G-I). Some of these *nubbin* mKO phenotypes displayed wild-type wing pattern elements (eyespots, submarginal bands) with only changes to pattern color (Supp. File 5). *nubbin* knockout also broadly affected scale morphology, producing wing scales that were shorter in length with rounder scalloped ridges (Fig. 3F-J) and affecting scale iridescence (Fig. 3F). Most *nubbin* mutants also were unable to properly unfold and expand their wings after eclosion.

**Fig. 3:**
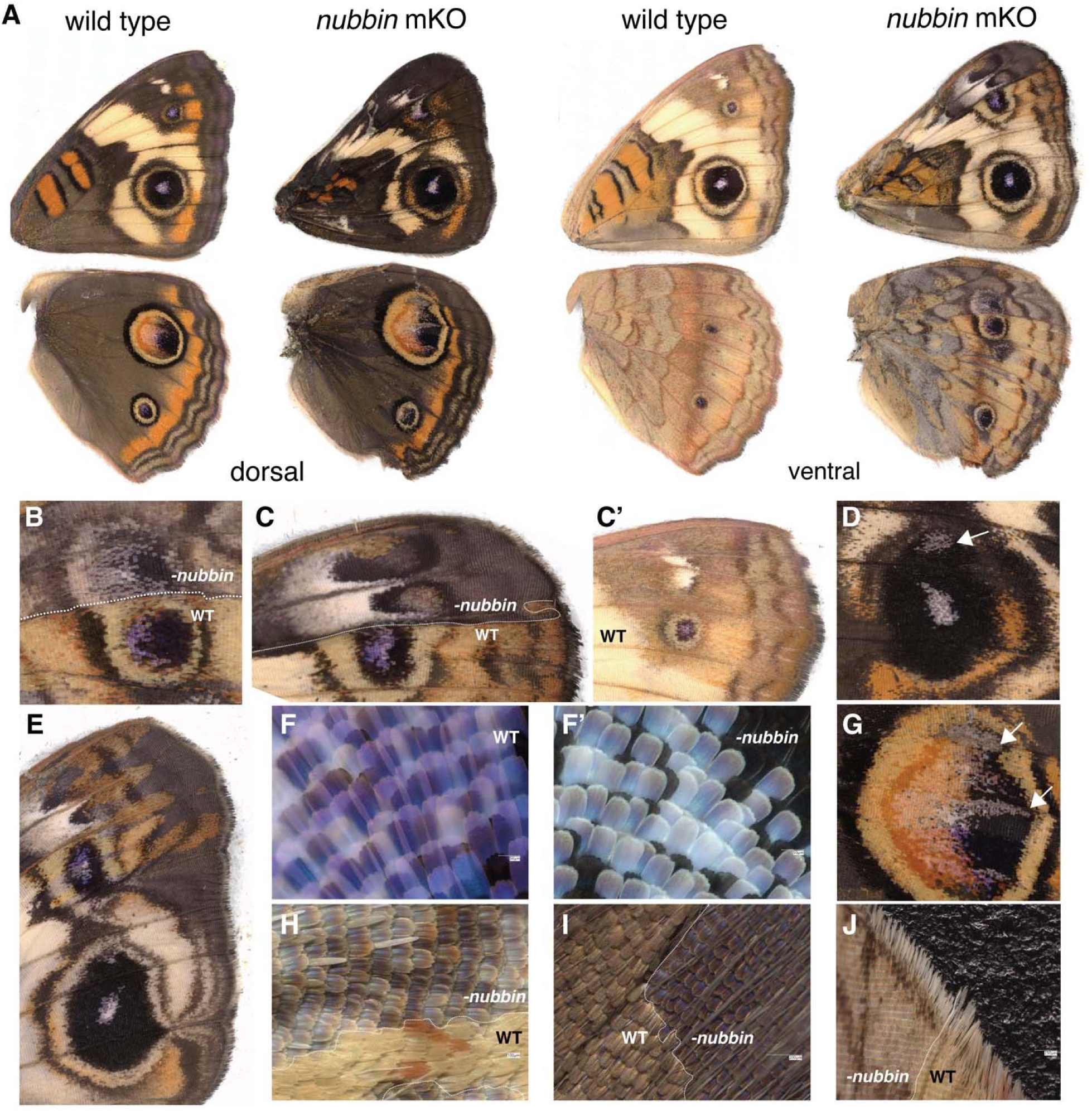
nubbin regulates wing patterning, pigmentation, and scale morphology. *nubbin* knockout broadly affected pattern development across the wing, including transformation of the border symmetry system, such as (B) induction of additional eyespots, (D) eyespot foci, and (C,E) larger eyespots, and (C,E) distortion of margin patterning. (B-C, E-J) *nubbin* knockout changed scale color identity, including loss of ommochrome pigmentation, as well as gain of black and gray melanin pigments. (F-J) *nubbin* knockout also affected wing scale morphology, (F-G) resulting in loss of iridescence and (F, H-J) resulting in wing scales that were shorter in length with rounder scalloped ridges compared to wild-type.

Some mKO individuals had changes in pattern formation in addition to pattern color, including aberrant eyespot number and size (Fig. 3B-E). Specifically, we observed ectopic eyespots within clonal regions (Fig. 3B), and these ectopic eyespots in large clonal regions were often larger than wild-type eyespots. We also observed additional eyespot foci, or eyespot centers, in otherwise wild type-like eyespots (Fig. 3D). These additional eyespot foci occurred at the midlines of vein compartments, corresponding to the initial *spalt* expression pattern during larval development (Fig. 2). Outside deletion clone regions, as inferred by reduced color pigmentation (see above), eyespots proximal to mKO wing veins were much larger in size (Fig. 3C, E). We also observed distortion of other elements of the border symmetry system. Wing margin patterns were stretched proximally, and sometimes there was fusion of the submarginal bands with the eyespots, often along the wing veins (Fig. 3B-C,E). The white anterior spots were also enlarged and stretched proximally (Fig. 3C,E). In sum, the overall effect was for color pattern elements to appear and/or expand in *nubbin* mKOs, leading us to infer that this gene plays a repressive role in some aspects of color pattern formation.

### 3.3 vvl regulates patterning of the wing veins and discal bands

Using in situ hybridization, we found *vvl* expressed in a proximal region of the wing disk during early fifth instar. *vvl* was also expressed along the boundaries of the developing lacunae throughout fifth instar, becoming restricted to only these developing wing veins later in development (Fig. 4). We did not detect expression of *vvl* at day 1, 2, or 3 of pupal wing development, although our RNA-seq data indicate that *vvl* was highly expressed later in day 3 and expressed moderately at day 7 of pupal development when we are no longer able to visualize mRNA expression using HCR due to autofluorescence (Supp. File 2).

**Fig. 4:**
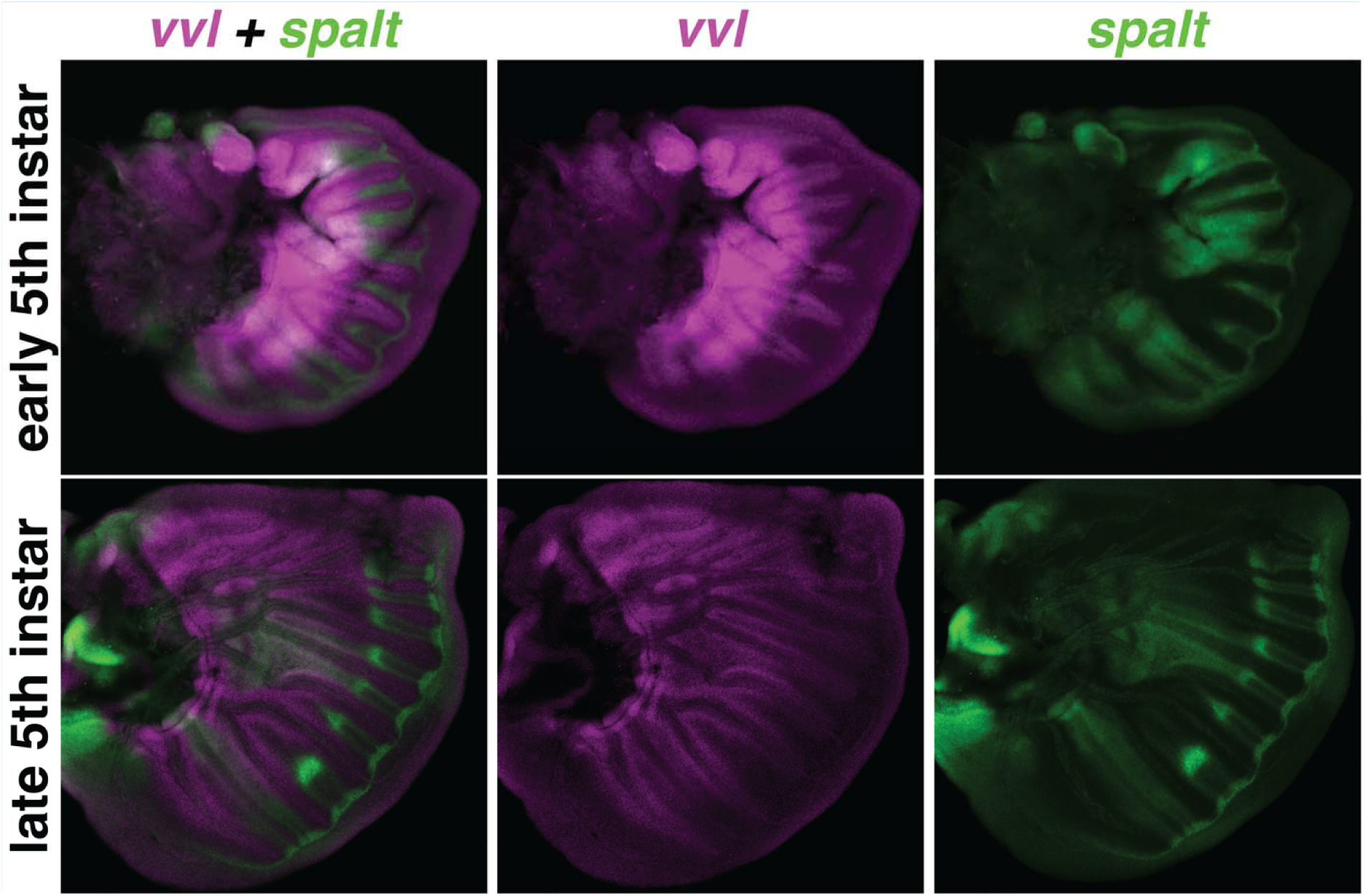
v*v*l is expressed in a proximal region of the wing during early fifth instar and the developing wing veins throughout fifth instar. *vvl* is expressed in a proximal region of the ring in early fifth instar wing disks. Throughout fifth instar, *vvl* is expressed along the boundaries of the developing lacunae.

CRISPR-Cas9 mKO of *vvl* induced ectopic veins (Fig. 5A), confirming that *vvl* plays a role in vein development. We also observed *vvl* mKOs with discal band color pattern aberrations, including loss of ommochrome pigmentation and expansion of melanin pigmentation (Fig. 5B), loss of both ommochrome and melanin pigmentation in the discal bands (Fig. 5C), and posterior expansion of a discal band (Fig. 5D), *vvl* knockout also resulted in upregulation of red pigmentation on the ventral hindwings (Fig. 5E).

**Fig. 5:**
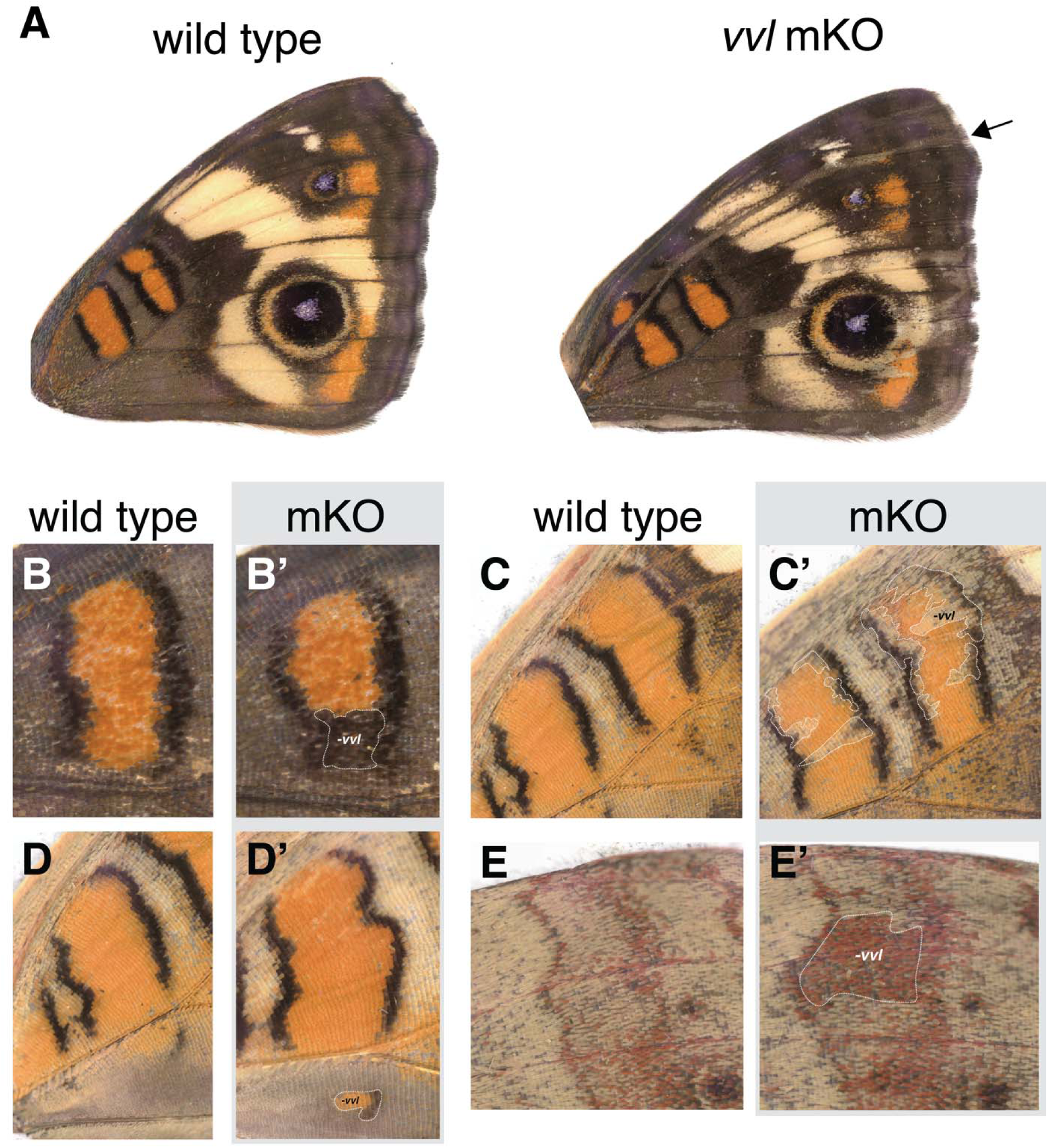
vvl determines vein and discal symmetry system patterning and ventral hindwing pigmentation. *vvl* knockout (A) induced an ectopic vein, (B) resulted in loss of ommochrome pigmentation of forewing discal bands, (C) loss of both ommochrome and melanin pigmentation in the ventral discal bands, (D) expansion of a ventral discal band, and (E) gain of ommochrome pigmentation on the ventral hindwing.

### 3.4 pdm3 regulates eyespot and wing margin patterning

In situ hybridizations revealed that *pdm3* has highly dynamic expression patterns over the course of wing development. In fifth instar wing disks we observed *pdm3* co-expressed with *spalt* in a band along the border lacuna (i.e. future wing margin). At day 1 of pupal development, *pdm3* was expressed in a wider proximal band along the border lacuna, while the expression domain of *spalt* extended further proximally from the wing margin compared to *pdm3*. At day 1, *pdm3* was also expressed in rings around the eyespot foci, as marked by *spalt* (Fig. 6). Expression of *spalt* and *pdm3* in the eyespot overlapped a few cells width at this stage. In day 2 pupal wings we observed *pdm3* co-expression with the lncRNA *ivory,* which marks the outer black eyespot ring (Fandino et al., 2024), and found that *pdm3* was co-expressed with *ivory* in a larger and thinner ring (Fig. 6). *pdm3* continued to be expressed in these eyespot rings during day 3 of pupal development (Supp. File 4, Fig. S1). These expression patterns suggest that *pdm3* may play a role in wing margin and eyespot patterning. Our RNA-seq analysis also found that *pdm3* was highly expressed toward the end of pupal development, at day 7 post-pupation, but HCR in butterfly wings is challenging at this stage due to autofluorescence of wing scales (Supp. File 2).

**Fig. 6:**
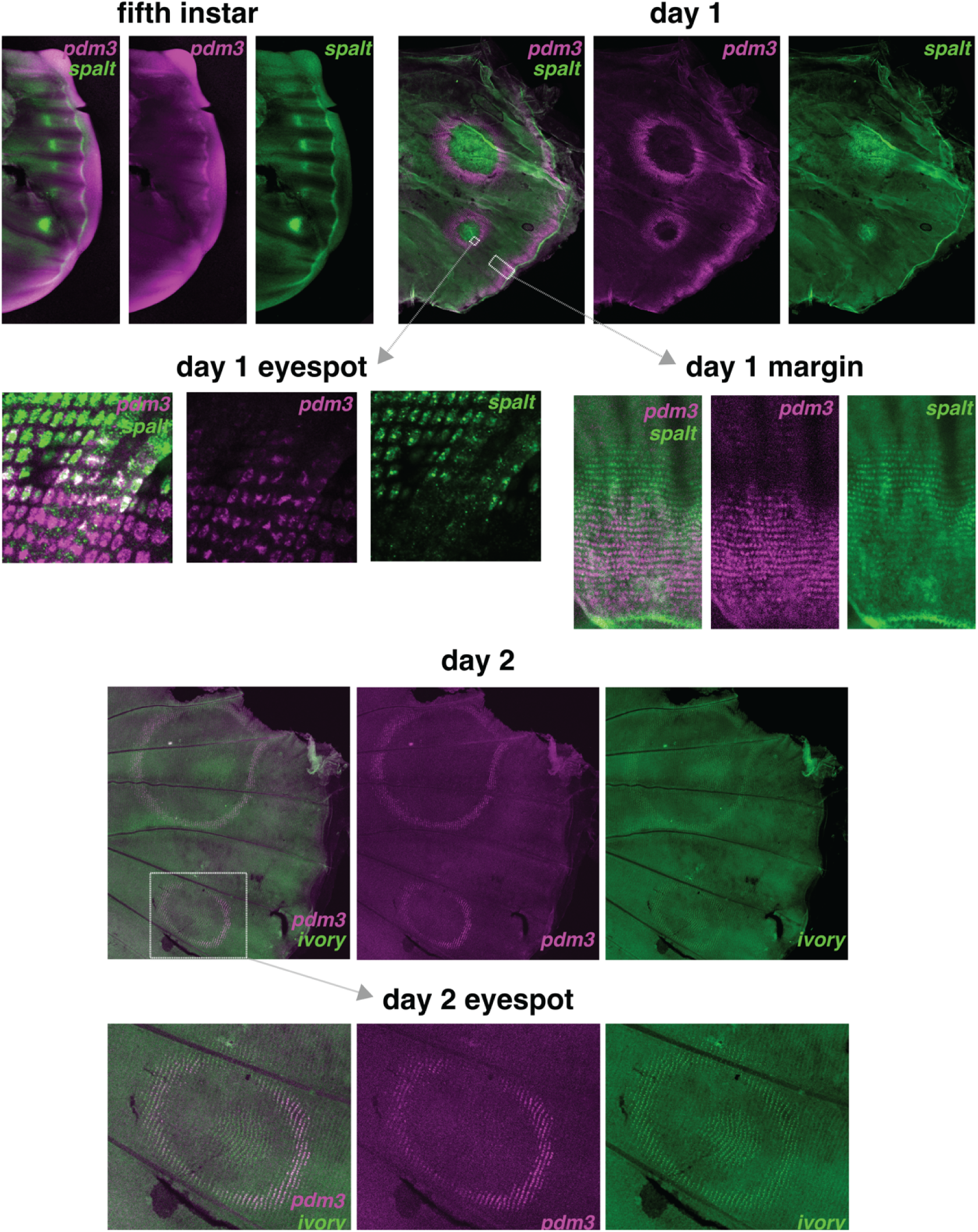
pdm3 is expressed in the future wing margins and eyespot rings. *pdm3* is co-expressed with *spalt* along the border lacuna during late 5th instar of larval development. At day 1 of pupal development, *pdm3* is expressed in an eyespot ring, which overlaps a few cells width with *spalt* expression in the center of the eyespot. *spalt* is co-expressed with *pdm3* in the wing margin, with *pdm3* expressed in a wider band from the border lacuna at day 1. *spalt* has a lower level of expression but a broader expression domain at the wing margin, expanding further proximally than *pdm3* from the border lacuna. In day 2 pupae, *pdm3* is co-expressed with the lncRNA *ivory* in a thinner and larger eyespot ring. *ivory* is also expressed in the center of the eyespot and along the border lacuna, but *pdm3* expression is no longer detectable in the wing margin at day 2.

CRISPR-Cas9 knockout of *pdm3* confirmed that this gene is necessary for multiple key aspects wing patterning – most noticeably in wing margin and eyespot pattern formation. Specifically, in *pdm3* mKOs we observed loss of both yellow and black outer rings of the eyespots (Fig. 7B-C, E-F, H-I), as well as fusion of the dark submarginal bands, which caused a loss of the light brown submarginal bands (Fig. 7D). On the dorsal wing surface, orange pigmentation also expanded from the wing margin and surrounded each eyespot (Fig. 7E, H-I). We also observed other effects of knocking out *pdm3* on wing color pattern, including gain of red and/or orange pigmentation across the distal region of the ventral wing (Fig. 7A, G, Supp. File 5) and loss of the black bars of the discal bands (Fig. 7J).

**Fig. 7:**
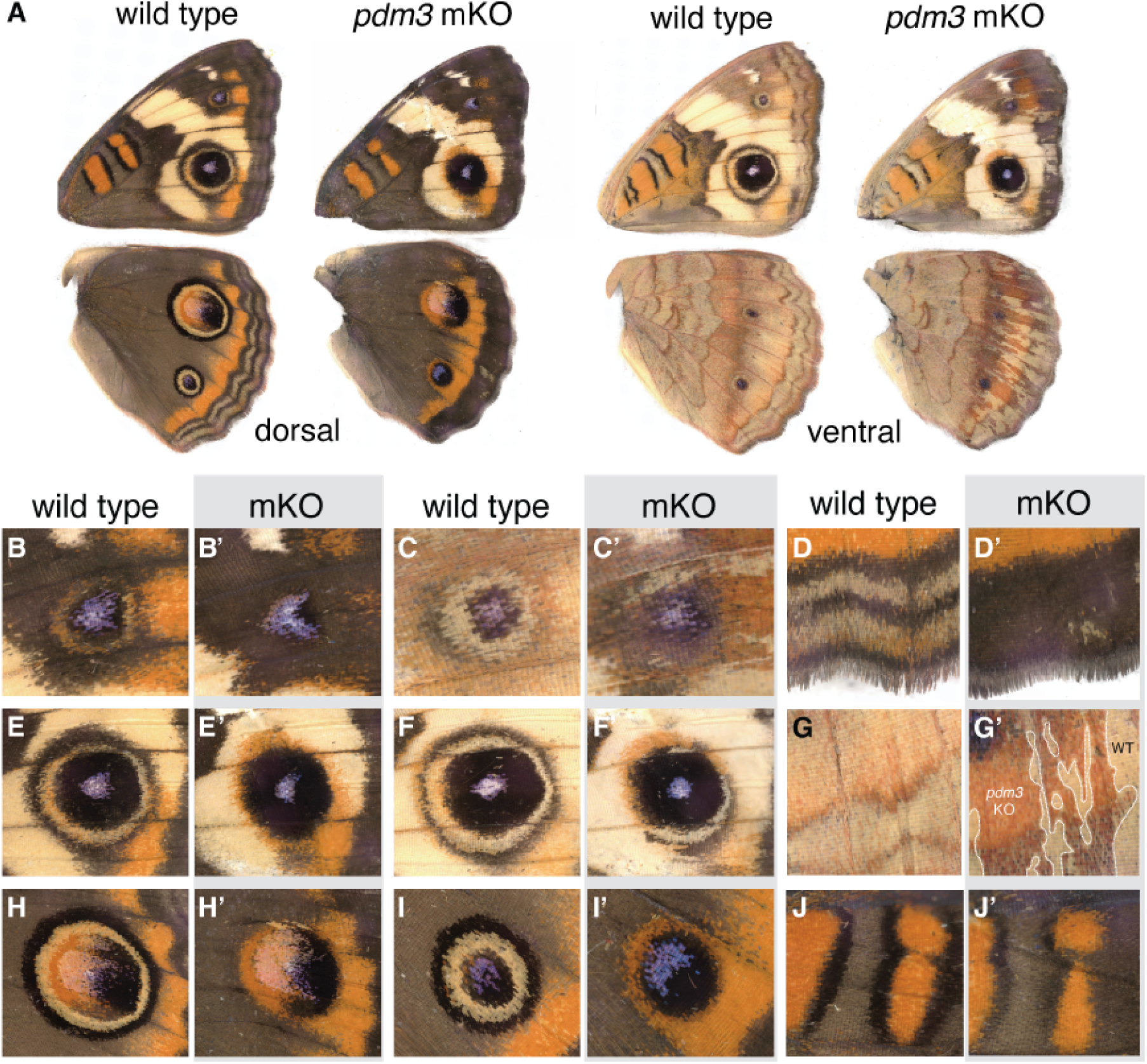
pdm3 regulates patterning of the discal bands and border symmetry system. *pdm3* mKOs resulted in (B-C, E-F, H-I) loss of all eyespot rings, (D) fusion of submarginal bands, (A,G) overall gain of ommochrome pigmentation in the distal region of the ventral wings, and (J) loss of black bars of the discal bands.

## 4. Discussion

### 4.1 nubbin knockout phenotypes are consistent with an eyespot repressing signal originating from wing veins

We observed that loss of *nubbin* resulted in ectopic eyespots and eyespot foci, as well as an increase in eyespot size (Fig. 3A-E). These effects on eyespot size and shape appeared to be non-cell autonomous since many of the effects occurred in eyespots outside of the deletion clones marked by pigmentation changes. We observed several mKOs with abnormally large eyespots that developed proximal to *nubbin* KO clone regions that included the nearest wing vein (Fig. 3C,E, Supp. File 5). *nubbin* was also expressed along the developing wing veins in fifth instar and pupal wing disks (Fig. 2). These phenotypes lead us to propose that *nubbin* regulates an intercellular repressor of eyespot determination originating at the developing wing veins.

The developing wing veins have long been suggested to play a repressive, compartmentalizing role in the eyespot determination process (Connahs et al., 2019; Nijhout, 1991; Schwanwitsch, 1924). The wing veins are the only differentiated structures at the time of eyespot specification (Nijhout, 1991). The center of each eyespot always develops at the midpoint between wing veins, and eyespot genes, such as *spalt*, are expressed at this midpoint in developing wing disks across nymphalids (Fig. 2), (Oliver et al., 2012; Reed et al., 2020; Reed and Serfas, 2004). Naturally occurring mutants with vein abnormalities also show changes in eyespot patterning. In *Bicyclus anynana,* for example, a single abnormally large eyespot develops across a much larger compartment that is missing a vein instead of two small eyespots that develop in two separate vein compartments in the wild-type (Brakefield et al., 1996). Similarly, incomplete development of a wing vein in a *Ypthima argus* mutant produced one large, merged eyespot with two foci instead of two distinct eyespots (Sekimura et al., 2015). These color patterns with improper vein development closely resemble color pattern abnormalities of *nubbin* mKOs (Fig. 3, Fig. 8), despite the fact the veins themselves remained intact. This leads us to infer that vein-associated signals, not the veins themselves, act as the pattern affecting agents.

**Fig. 8:**
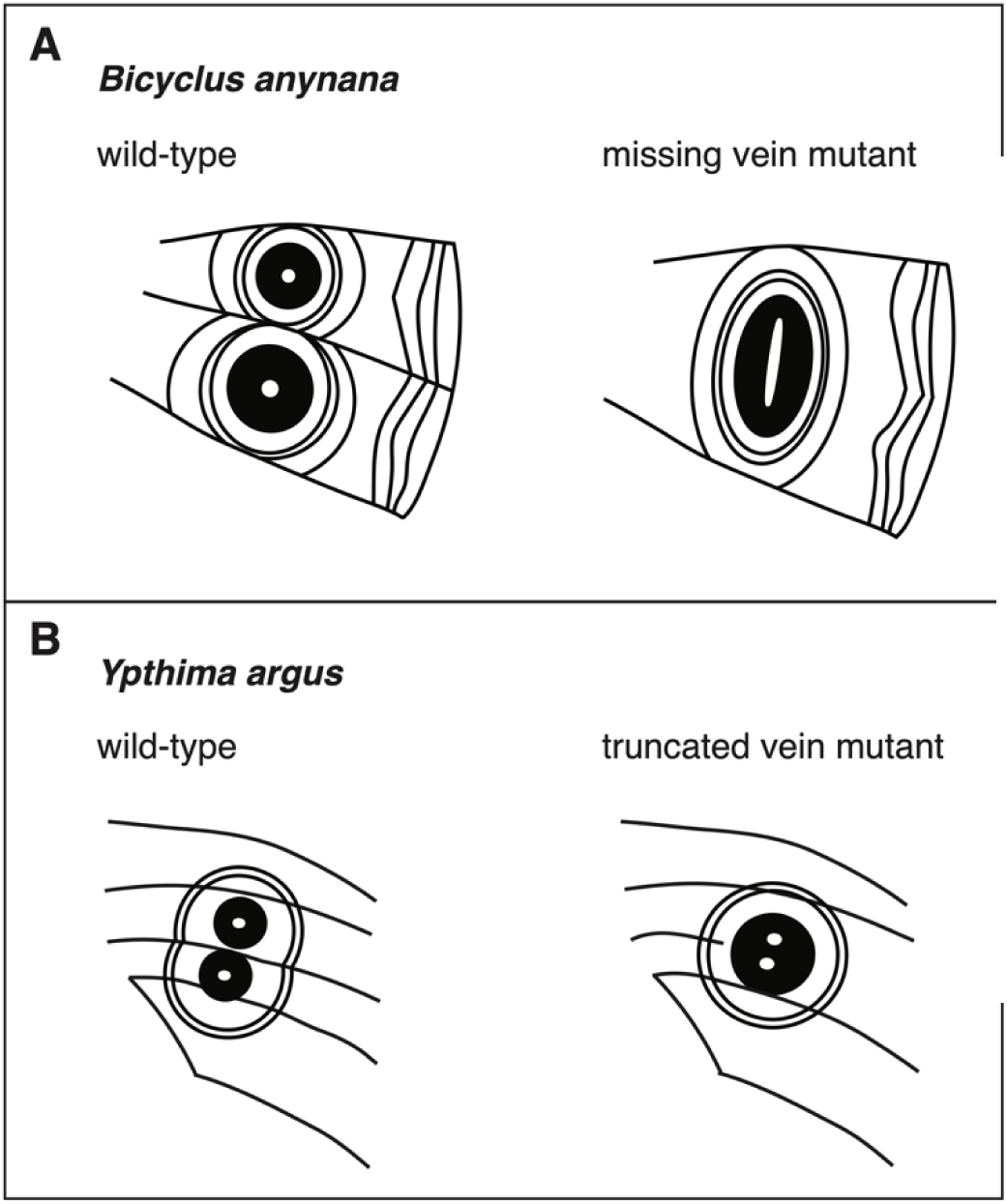
Disruption of wing vein development affects eyespot patterning in spontaneously-occurring mutants. (A) The *Bicyclus anynana* mutant *Cyclops* is missing a vein, resulting in one large eyespot in place of two eyespots on the ventral hindwing (Brakefield et al., 1996). (B) A *Ypthima argus* mutant with incomplete vein development has a single eyespot with two foci in place of two eyespots on the ventral hindwing (Sekimura et al., 2015).

We propose the most parsimonious model to explain our data is one where *nubbin* regulates an intercellular repressor at the wing veins, which then diffuses and decays with increasing distance from the veins toward the intervein midlines (Fig. 9). We hypothesize that this signal represses the determination of the eyespot focus. The eyespot focus produces a morphogen during early pupal development that diffuses outward and induces eyespot pattern development (Nijhout, 1980). We suggest that loss of this repressor in a *nubbin* KO wing vein results in a larger eyespot (as in Fig. 3C) because more cells differentiate into focal eyespot cells, which then release a stronger morphogen gradient across a larger region of the wing (Fig. 9).

**Fig. 9:**
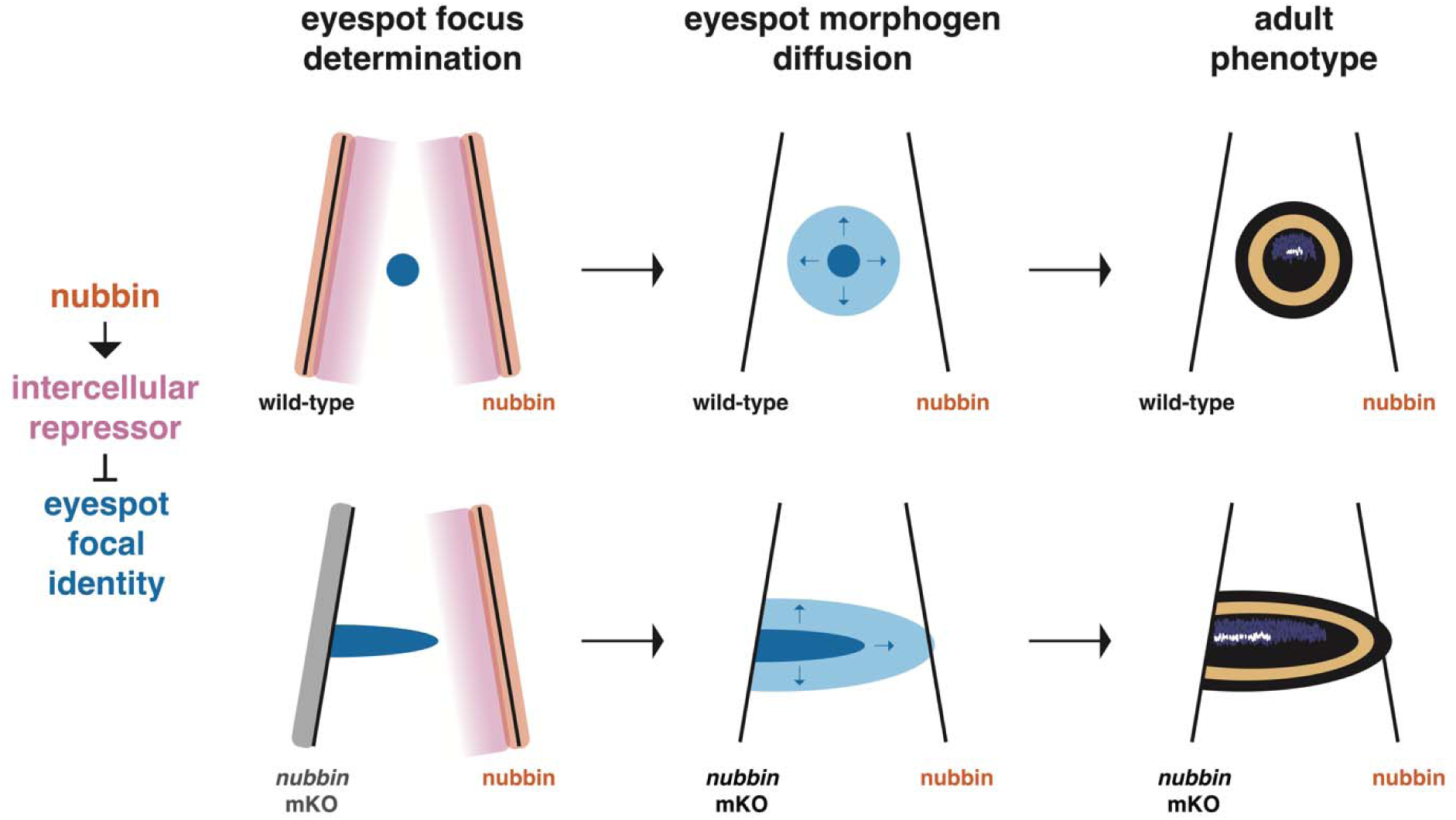
Model of nubbin regulation of eyespot development via an intercellular repressor of eyespot focal determination. An increase in eyespot size due to loss of *nubbin* expression in a proximal wing vein, as in the *nubbin* mKO in Fig. 3C, can be explained by a simple model where *nubbin* upregulates an intercellular repressor at the developing wing vein boundaries. This repressor diffuses from the veins toward the midline and suppresses cells from adopting eyespot focal identity. Eyespot focal cells then produce the eyespot morphogen, which diffuses outward (for large eyespots, this domain can extend beyond the wing veins) and induces eyespot pattern development. Cells near the *nubbin* mKO vein are not suppressed from developing into eyespot focal cells as they are in wild-type, resulting in a larger eyespot focus, higher morphogen levels, and a larger eyespot pattern on the adult wing.

**Fig. 10:**
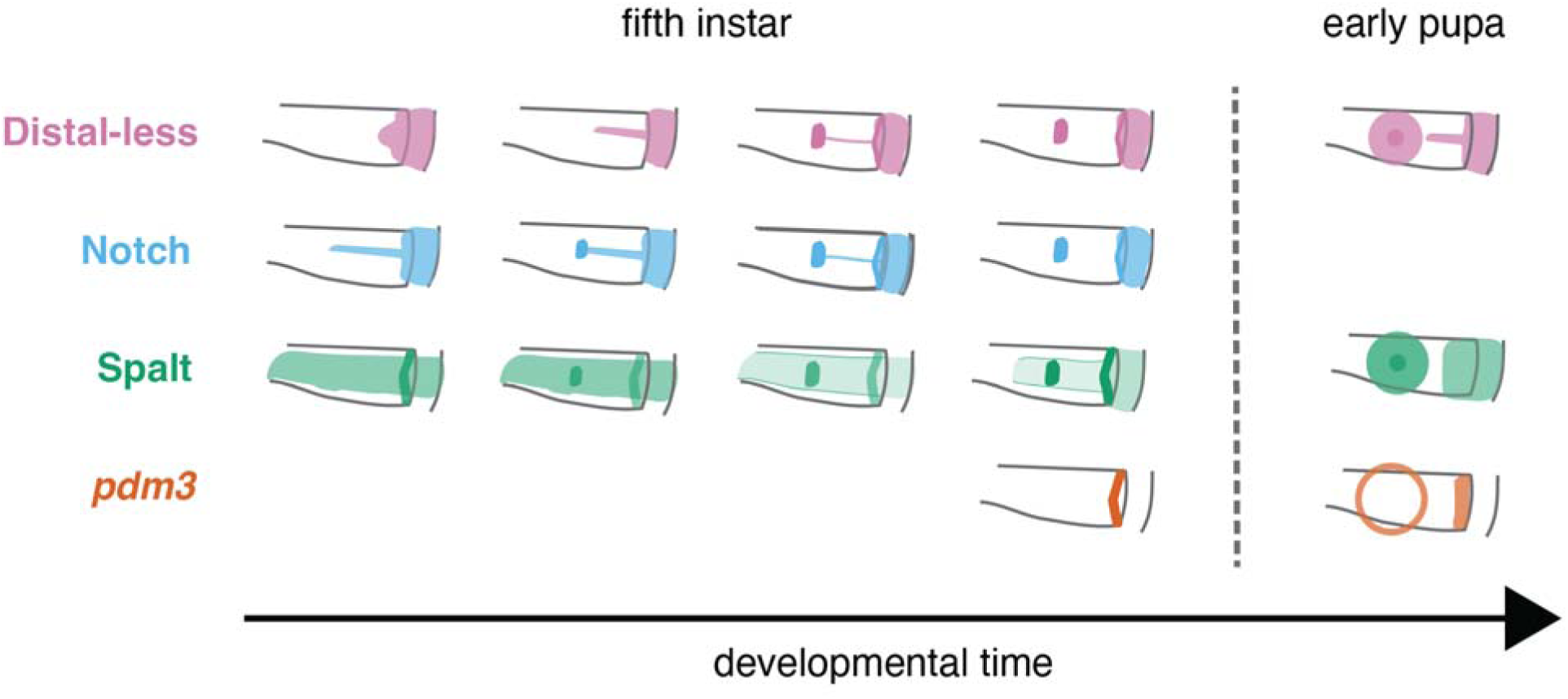
Spatiotemporal expression patterns of shared eyespot and wing margin genes. In early fifth instar larval wing disks, Distal-less, Notch, and Spalt proteins are all expressed in the wing margin; Spalt is expressed along the midline of the vein compartment, and Notch is expressed in a narrower, shorter projection from the margin along the midline. As the wing disk develops, Distal-less expression begins to expand in a similar projection to Notch from the wing margin along the midline. Ultimately, Distal-less, Notch, and Spalt are all specifically upregulated in the eyespot and wing margin patterns by late fifth instar. *pdm3* is only expressed in the wing margin during late fifth instar. At day 1 of pupal development, Distal-less, Spalt, and *pdm3* are co-expressed in the eyespot and wing margin. Expression pattern illustrations of a representative vein compartment of the wing were based on Fig. 4, Fig. 6 and (Brunetti et al., 2001; Reed and Serfas, 2004; Wee et al., 2022), which characterized these patterns using immunohistochemistry (Distal-less, Notch, Spalt) and in situ hybridization (*pdm3*) across multiple nymphalid species, including *B. anynana* and *J. coenia*.

Similarly, we propose that ectopic eyespots and eyespot foci in *nubbin* mKOs are also due to downregulation of this intercellular signal originating from the wing veins (Fig. 3B,D). Interestingly, in *J. coenia* and many other nymphalid species, most vein compartments express *spalt* and other eyespot genes in a spot pattern at the fifth instar stage (Fig. 2) (Oliver et al., 2012; Reed et al., 2020, 2007). However, most of these spots will not give rise to eyespots in the adult wing. In fact, by early pupal development, *spalt* is only expressed in two spots that correspond to the adult eyespots in *J. coenia*, and only the centers of these vein compartments are able to induce eyespot patterning when transplanted (Fig. 2, Nijhout, 1980). Our data suggest that all eyespot foci specified at fifth instar have the potential to generate mature eyespot foci, but some of these potential eyespots are completely inhibited by a *nubbin*-activated signal.

Both the number and size of eyespots have evolved many times throughout nymphalid evolution (Ho et al., 2016; Nijhout, 1991). The size and number of eyespots can influence predation (Chan et al., 2021; Ho et al., 2016; Lyytinen et al., 2004), and eyespot size and UV iridescence are also sexually-selected traits in some species (Robertson and Monteiro, 2005). We speculate that differences between whether a vein compartment develops an eyespot or not, as well as relative size differences between eyespots of different compartments, could be determined by regional differences in the expression of some co-regulator(s) of the intercellular repressor. These co-regulator(s) could influence either: a) the production of this repressor, b) the rate of repressor degradation, or c) the sensitivity of cells to repression. Therefore, changes in expression of the co-regulator(s) could explain frequent evolutionary changes in eyespot number and size since both the eyespot prepattern (Oliver et al., 2012) and the capacity to induce the downstream eyespot developmental program are conserved across vein compartments. This idea could be tested after these co-regulators and the vein-originating repressor are identified, such as through a large-scale screen of wing-expressed genes.

Knockout of *nubbin* also transformed other pattern elements specified by *spalt*, including the wing margin bands and the anterior white spots of the forewing. In *nubbin* mKOs, these elements were stretched proximally compared to wild-type, changing both their relative positions and sizes on the wing (Fig. 3C,E). Therefore, we can infer that *nubbin* also regulates the position and shape of other elements of the border symmetry system. We cannot conclude from these data alone whether *nubbin* regulates these patterns via the same mechanism coordinating eyespot development. However, it would not be surprising for regulation of wing margin patterns to occur via a signal originating at the wing vein. In *J. coenia,* submarginal bands generally form triangles or chevrons within each vein compartment, with the most proximal point at the midline between the veins. The wing veins of many lepidopteran species demarcate repeated patterns that are symmetrical across the midline of each compartment (for example see the moths *Rheumaptera undulata* and *Habrosyne scripta*), suggesting some intercellular signaling from the wing veins could be at play here. An exciting open question is whether a deeply conserved mechanism coordinates the early positioning of these patterns within vein compartments that has been abundantly elaborated upon to generate a diversity of adult color patterns, or if the wing veins have repeatedly evolved mechanisms to coordinate color pattern across many different taxa.

### 4.2 nubbin controls scale color and morphology

We also found that *nubbin* regulates the pigmentation and morphology of scales across the wing. *nubbin* mKOs developed wing patterns in grayscale rather than in color. Scales that produce ommochromes in wild-type were depigmented in mKOs, and scales that produce presumptive brown melanins in wild-type instead produced gray melanins in mKOs (Fig. 3). We also observed that some mKOs displayed differences in scale color identity and morphology without differences in color pattern from wild-type, suggesting that *nubbin* regulates scale color and morphology later in development than pattern formation, such as during day 3 of pupal development when *nubbin* is expressed ubiquitously (Fig. 2) or later in development (Supp. File 2). These observations suggest that *nubbin* is required for ommochrome synthesis and modifies the chemical synthesis of melanins. This loss of ommochrome pigmentation in *nubbin* mKOs resembles the loss of ommochrome pigmentation across the wing in *optix* mKO butterfly wings; however, *nubbin* mKOs differ from *optix* mKOs in other effects on pattern, pigmentation, and scale morphology (Zhang et al., 2017). This suggests that these two factors could be in a shared ommochrome pathway, while also regulating other wing color pattern genes independently of each other. More work is needed to determine the relationship between these genes and the downstream patterning network.

We also observed differences in scale morphology across the wing in *nubbin* mKOs versus wild-type: *nubbin*-deficient scales were consistently shorter in length with rounder scalloped ridges (Fig. 3F,H-J). *nubbin* is not the first gene to be found to regulate both scale morphology and color; other genes, including *optix* and *yellow*, also affect both traits across the butterfly wing (Matsuoka and Monteiro, 2018; Zhang et al., 2017). Scanning electron microscopy or helium ion microscopy could reveal more detail about the structural differences in *nubbin* mKO scales.

### 4.3 vvl regulates vein development and color pattern readout

Comstock and Needham (1910) first inferred homology of wing venation patterns across insects over a hundred years ago (De Celis and Diaz-Benjumea, 2003), yet we still do not know to what extent the developmental process of venation is conserved. Vein development has primarily been studied in *D. melanogaster,* and only a few conserved venation factors have been identified in other insects, such as *Dpp* in the sawfly and *spalt* in butterfly wings (Banerjee and Monteiro, 2025, 2020; Matsuda et al., 2013; Shimmi et al., 2014; Zhang and Reed, 2016).

Here, we found that *vvl* regulates vein development and that it has undergone a heterochronic shift in expression in the butterfly wing. We observed *vvl* expression across the proximal region of the wing and along the developing wing veins throughout the last instar in *J. coenia*. *vvl* expression was restricted to only the developing wing veins late in the last instar (Fig. 4). In contrast, during the last instar of *D. melanogaster, vvl* is expressed in multiple wing regions, throughout both wing blade and hinge, and disappears from the intervein regions during early pupal development (de Celis et al., 1995). These early pupal fly wing disks then express *vvl* along their developing wing veins in a similar pattern to last instar *J. coenia* wing disks.

The effects of *vvl* mutation also suggest that this gene plays a role in vein development, but it is not clear whether this is a conserved function with *Drosophila*. We found that knockout of *vvl* results in ectopic wing vein development in *J. coenia* (Fig. 5A), whereas *D. melanogaster vvl* mutant clones either fail to develop wing veins or develop thicker veins than wild-type (de de Celis et al., 1995). Given that these mutant phenotypes are from mosaic mutants, more work is needed to ascertain whether these are due to differences in *vvl* function between the two insects or differences in the timing and/or location of mutations induced across the wing.

We also found that *vvl* has evolved a novel role in color pattern regulation in butterflies compared to flies. *vvl* had previously been associated with color pattern variation in the distal region of the forewing in *Heliconius* butterflies but had not been functionally tested (Van Belleghem et al., 2017). Surprisingly, our knockout phenotypes do not correspond to the distal forewing. Instead, we observed *vvl* knockout causes both gain and loss of ommochrome and melanin pigmentation in the discal bands (Fig. 5B-D). Therefore, we infer that *vvl* is involved in multiple stages of patterning readout. We also found that there was a gain of red pigmentation in the ventral hindwing, which could also be the result of *vvl* regulating patterning readout in this region of the wing (Fig. 5E). Again, due to mosaicism, we cannot rule out that *vvl* has a function that better corresponds to previous predictions from mapping experiments in *Heliconius*. We can only conclude that *vvl* plays an unexpected role in the pigmentation of the discal bands and ventral hindwing.

### 4.4 pdm3 coordinates development of outer eyespot rings and wing margin bands

Our CRISPR knockouts reveal that *pdm3* is necessary for patterning the outer yellow and black rings of all eyespots in *J. coenia* (Fig. 7A-C, E-F, H-I). These results are consistent with the loss of outer yellow and black eyespot rings observed in *V. cardui* CRISPR knockouts (Loh et al., 2025a), suggesting that this is a conserved role for *pdm3* in nymphalid butterflies.

Our in situ hybridization data lead us to speculate that *pdm3* expression is responding to the eyespot morphogen. At day 1 of pupal development, *pdm3* is expressed in a thicker ring with a smaller circumference. This expression overlaps a few cells width with *spalt,* which is expressed in the center of the eyespot pattern during early pupal development (Fig. 6). At day 2 of pupal development, we found that *pdm3* is expressed in a thinner ring with a much larger circumference. This expression domain corresponds directly with the expression of the lncRNA *ivory* in this outer ring at day 2 of pupal development (Fig. 6). Previous research has shown that *ivory* is the host gene for the microRNA *mir-193* and is necessary for development of the outer melanic eyespot ring in *V. cardui, B. anynana,* and *J. coenia* (Fandino et al., 2024; Hanly et al., 2026; Livraghi et al., 2024; Tian et al., 2024). We propose that this spatiotemporal pattern of *pdm3* expression could reflect a response to spatiotemporal changes in the concentration of a diffusing morphogen. While an eyespot morphogen has been inferred from transplant experiments, we do not know its identity nor the dynamics of its release. The pattern of *pdm3* expression that we observed could be explained by several different processes of morphogen readout. One possible mechanism is that a long-range morphogen, produced at a constant rate, could increase in concentration over time across the eyespot rings, such that the morphogen concentration for activating *pdm3* expression is initially present proximal to the eyespot center, or the proposed source of the morphogen, and later, is present further away from the center. Under this model, *pdm3* expression is suppressed by higher concentrations of morphogen. Another possible explanation is that the eyespot rings could be induced by sequential short-range morphogens, i.e. a secondary morphogen could be responsible for inducing the day 2 *pdm3* expression pattern. We propose that within the first 8-24 hours of pupal development, when *pdm3* is expressed in a smaller circumference ring, *pdm3* regulates yellow ring differentiation. During the second day of pupal development, when *pdm3* is expressed in the larger circumference ring, we suspect that *pdm3* activates genes responsible for patterning the black ring, such as *ivory.* Further investigation into the eyespot patterning network would help clarify this.

Our CRISPR knockouts also revealed that *pdm3* regulates the development of the wing margin bands. The submarginal bands appeared to have fused and formed a single, larger marginal band of dark brown on both forewing and hindwing in *pdm3* mKOs, resulting in a loss of the light tan bands. Further, we observed that *pdm3* was co-expressed with *spalt* in a thin, chevron-like band near the border lacuna at fifth instar, and later, its expression domain expanded proximally during early pupal development. Future work could determine to what degree the network downstream of *pdm3* is shared between the wing margin and eyespot.

### 4.5 Proposed homology of eyespot and wing margin gene regulatory networks

Our observation that *pdm3* is co-expressed with *spalt* in both the eyespot and wing margin (Fig. 6), when considered with prior work on the expression of other shared genes in these traits, supports the hypothesis that the eyespot evolved in nymphalids via co-option of the wing margin patterning network (Held, 2013). Four key developmental factors, *pdm3,* Spalt, Distal-less (Dll), and Notch, all show a pattern of expression where they are initially more strongly expressed in the wing margin, and later are specifically upregulated in the eyespot and wing margin patterns (Fig. 6, 10) (Carroll et al., 1994; Reed and Serfas, 2004). Interestingly, Dll and Notch expression are characterized by projections from the wing margin that extend along the midline of the vein compartment before eyespot specification. Spalt is expressed along the midline throughout the vein compartment (Fig. 4, 10). In outgroup species without eyespots, there is also conserved expression in the wing margin of Dll, Notch, and Spalt, and conserved midline projections of Dll and Notch (Reed and Serfas, 2004; Shirai et al., 2012), indicating these genes’ roles in patterning the wing margin evolved before the eyespot and are deeply conserved in Lepidoptera.

There is also functional evidence to support this idea: *pdm3, spalt,* and *Dll* are all required for proper development of the wing margin bands and eyespot patterns (Fig. 7, Zhang and Reed, 2016). Further, in our *nubbin* knockouts, both the eyespot and wing margin patterns increased in size and were shifted proximally, which indicates a shared upstream factor in the regulation of these traits (Fig. 3).

While co-option of a gene regulatory network is a popular hypothesis for the origin of the eyespot, many other potential ancestral networks have been proposed, including the antenna, leg, wing vein, and wound-healing networks (Banerjee and Monteiro, 2025; Monteiro, 2015; Murugesan et al., 2022; Özsu and Monteiro, 2017). More detailed dissection of these networks, including the specific regulatory interactions of these and other genes, is needed to determine which network, if any, was co-opted for eyespot patterning.

### 4.7 Conclusion

Overall, we found that a clade of POU domain factors has played an important role in the evolution of butterfly wing color pattern. All three of the factors we investigated play multiple, major roles in wing patterning and pigmentation. We also found new leads for resolving persistent mysteries in butterfly wing pattern development. For example, *nubbin* mosaic knockouts provided the first experimental evidence for a century-old hypothesis that the wing veins position and regulate the development of the butterfly eyespot. Another striking finding was the dynamic expression of *pdm3* in the wing margin followed by expression in the eyespot, which we found was similar to the spatiotemporal pattern of other eyespot genes. Our knockout data confirmed that *pdm3* regulates both eyespot and wing margin development, which suggests that the eyespot may have evolved via co-option of the wing margin patterning network, and future work can explore the relative homology of these patterning mechanisms in more depth. These data further illustrate that individual genes with ancestral roles in insect wing development, such as *vvl* and *nubbin,* have been co-opted for new roles in butterfly color pattern, a long-running theme of evo-devo. By characterizing the role of *nubbin, vvl,* and *pdm3,* we aim to contribute to the long-term project of describing the color pattern formation process across butterfly species to better understand how the developmental programs of morphologies evolve.

## CRediT authorship contribution statement

**Jeanne M. C. McDonald** – Conceptualization, Investigation, Funding acquisition, Data curation, Validation, Supervision, Writing – original draft, Writing – review & editing. **Qin Guo** – Investigation, Writing – review & editing. **Seydeanna Delgado** – Investigation, Writing – original draft. **Connor A. Amendola** – Investigation, Writing – review & editing. I**ya A. Garg** – Investigation. **Robert D. Reed** – Conceptualization, Funding acquisition, Supervision, Writing – review & editing.

## Appendix A. Supplementary data

Supplementary File 1. – Phylogenetic analysis

Maximum likelihood tree for nubbin, vvl, and pdm3 orthologs across Pancrustacea and in a mollusc species, in addition to holometabolous insects in Fig. 1, confirmed inferred orthology and expected phylogenetic relationships.

Supplementary File 2. – Gene expression data

Normalized expression data from RNA-seq across developmental stages in the wing of *Junonia coenia* for *nubbin, vvl,* and *pdm3*.

Supplementary File 3. – Genotyping results

Inference of CRISPR Edits analysis results for each mKO genotyped.

Supplementary File 4. – Additional HCR for *pdm3*

HCR for *pdm3* at day 3 of pupal development, z-stack for *pdm3* and *spalt* at day 1 of pupal development, and z-stack for *nubbin* at day 3 of pupal development..

Supplementary File 5. – Additional images of mKOs

Additional images of both left and right wings from mKOs, including identification numbers for each mKO.

Table S1. – Summary of mKO phenotypes

Excel sheet summarizing the mutant phenotypes for each gene, including the corresponding identification numbers for each mKO as reported in Supp. File 5.

## Acknowledgements

The authors would like to thank Jillian Taveira, Nigel Williams, and Isabella Toso for assistance with maintaining the lab population of *J. coenia*. We thank members of the Reed Lab, Babonis Lab, Jason Dombroskie, Kevin McDonald, and Cece Measán for helpful discussion. We thank Johanna Dela Cruz and Rebecca Williams for assistance with imaging.

## Data availability

All code and protein sequences are available at: https://github.com/jeannexpression/pdm_tree

