## Supplemental File 1 for "*nubbin*, *ventral veinless*, and *pdm3* play diverse roles in butterfly wing pattern development"

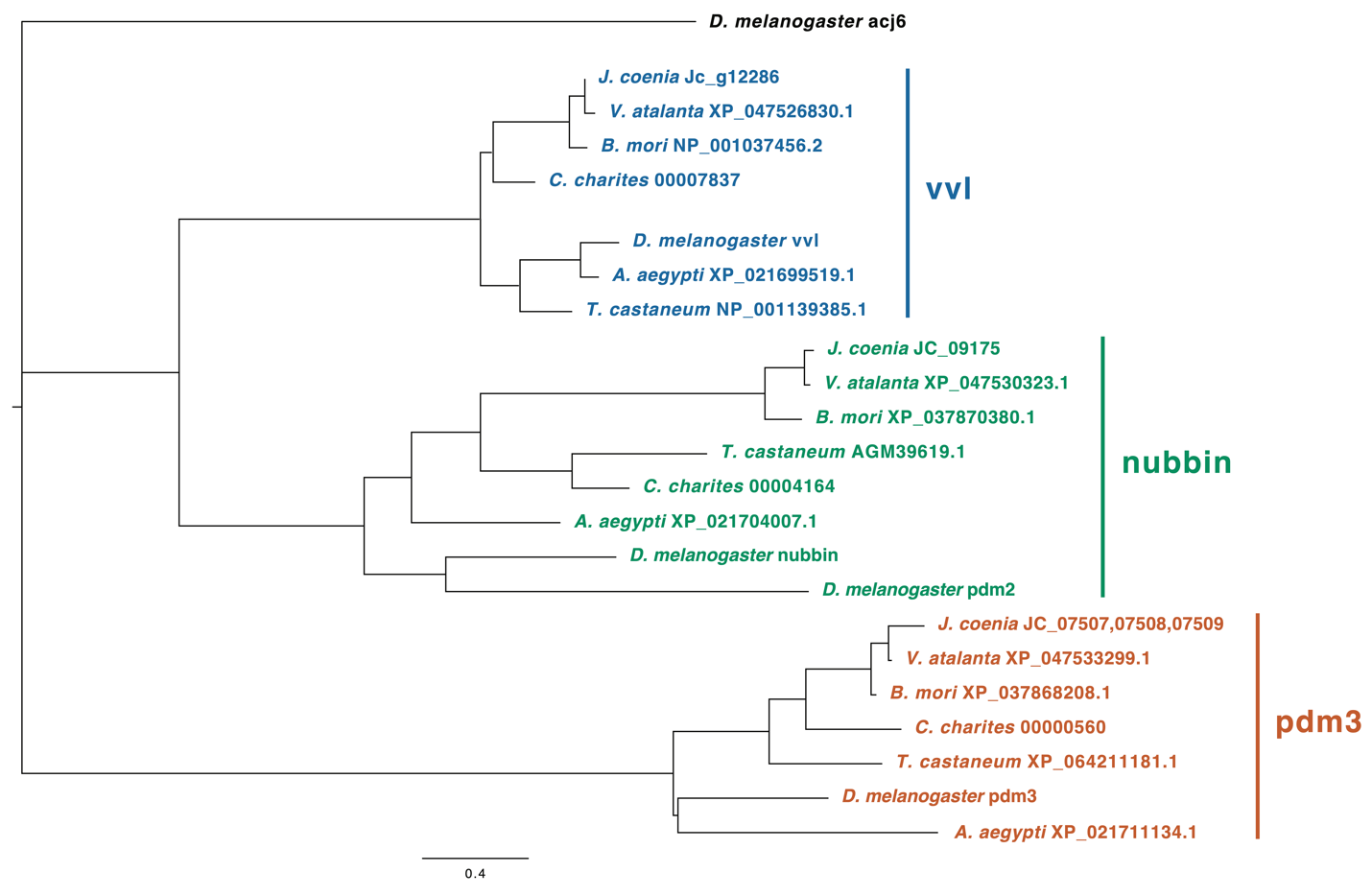

**Fig. S1: Maximum likelihood tree for vvl, nubbin, and pdm3 protein sequences.**

Phylogenetic relationships of vvl, nubbin, and pdm3 orthologs across insects inferred from a maximum likelihood consensus tree. *D. melanogaster* acj6, a more distantly related POU domain factor, is included as an outgroup. The scale bar represents the number of substitutions per site.

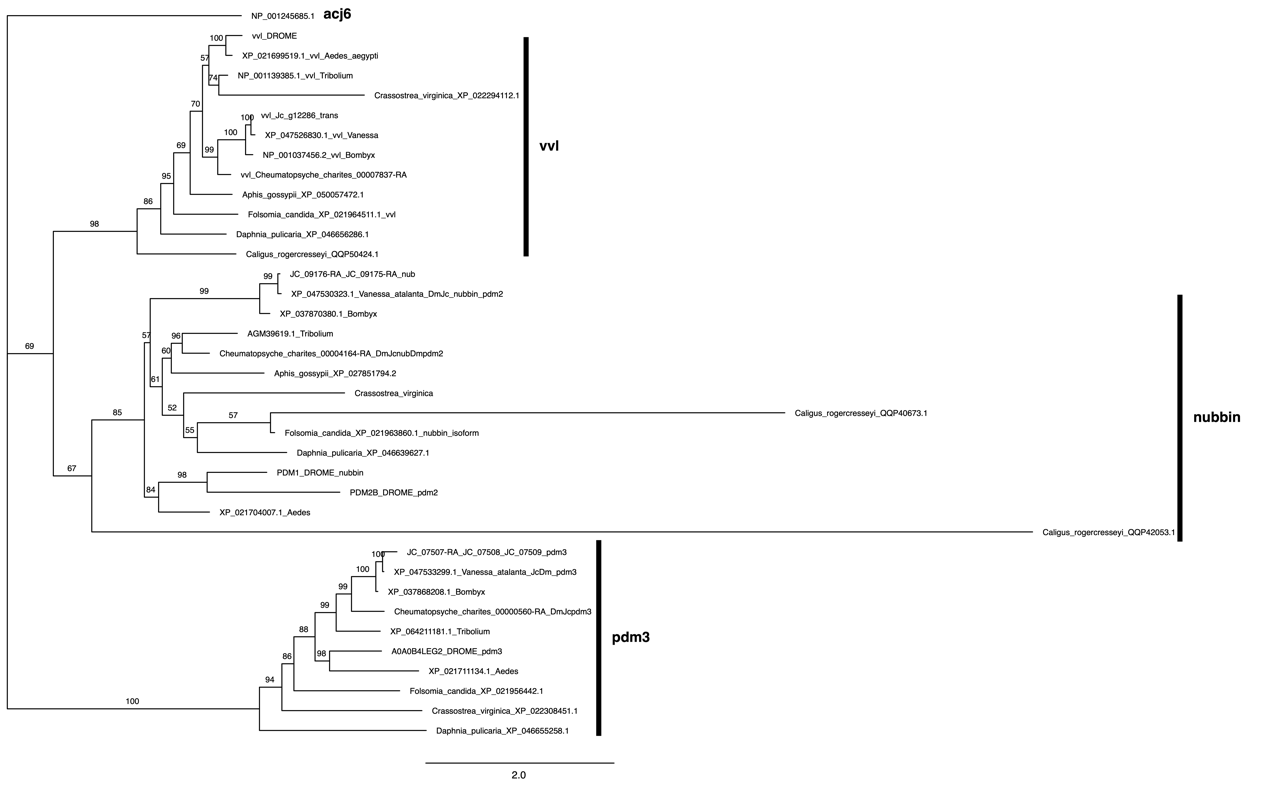

### **Fig. S2:** **Maximum likelihood tree for vvl, nubbin, and pdm3 protein sequences with additional species.** Phylogenetic relationships of vvl, nubbin, and pdm3 orthologs across Pancrustacea and a mollusc outgroup were inferred from a maximum likelihood consensus tree constructed from 1000 bootstraps and were consistent with our tree across holometabolous insects (Fig. 1). *D. melanogaster* acj6, another POU domain factor, is included as an outgroup. The scale bar represents the number of substitutions per site. Numbers in parentheses are bootstrap supports (%).
