## Supplemental File 2 for "*nubbin*, *ventral veinless*, and *pdm3* play diverse roles in butterfly wing pattern development"

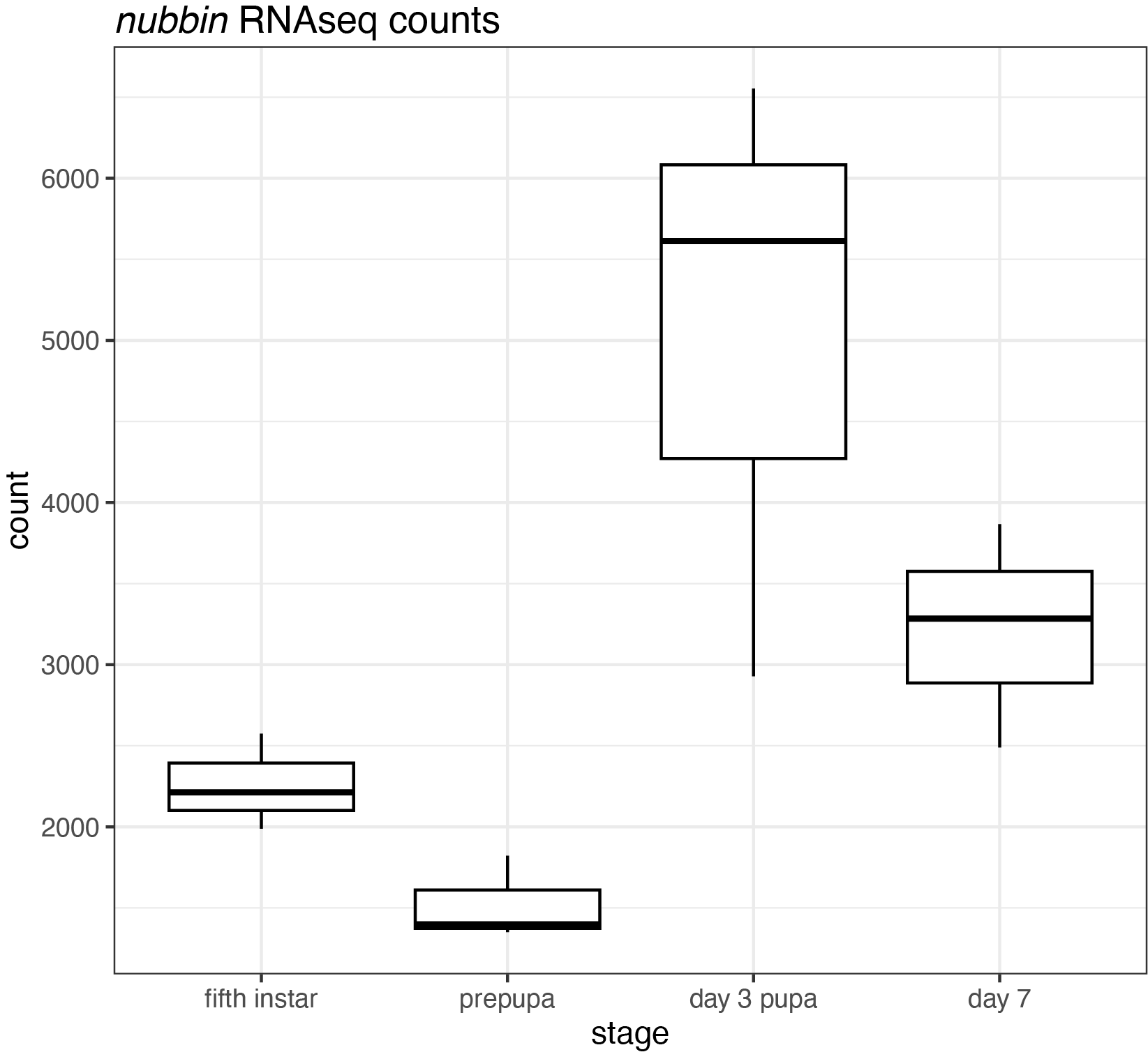

**Fig. S3: Normalized counts of *nubbin* transcripts in the wing across development in *Junonia coenia.***

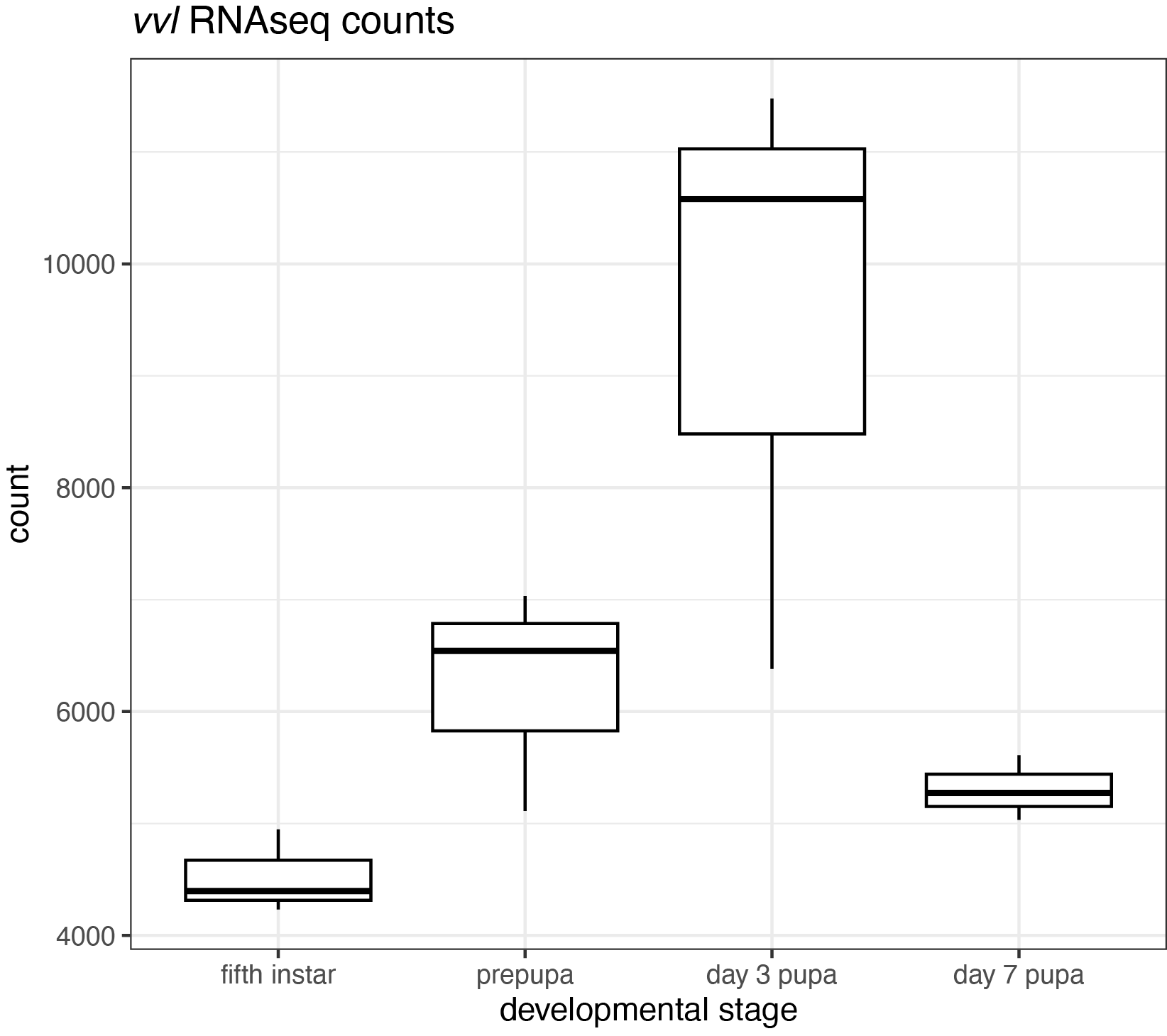

**Fig. S4: Normalized counts of *vvl* transcripts in the wing across development in *Junonia coenia.***

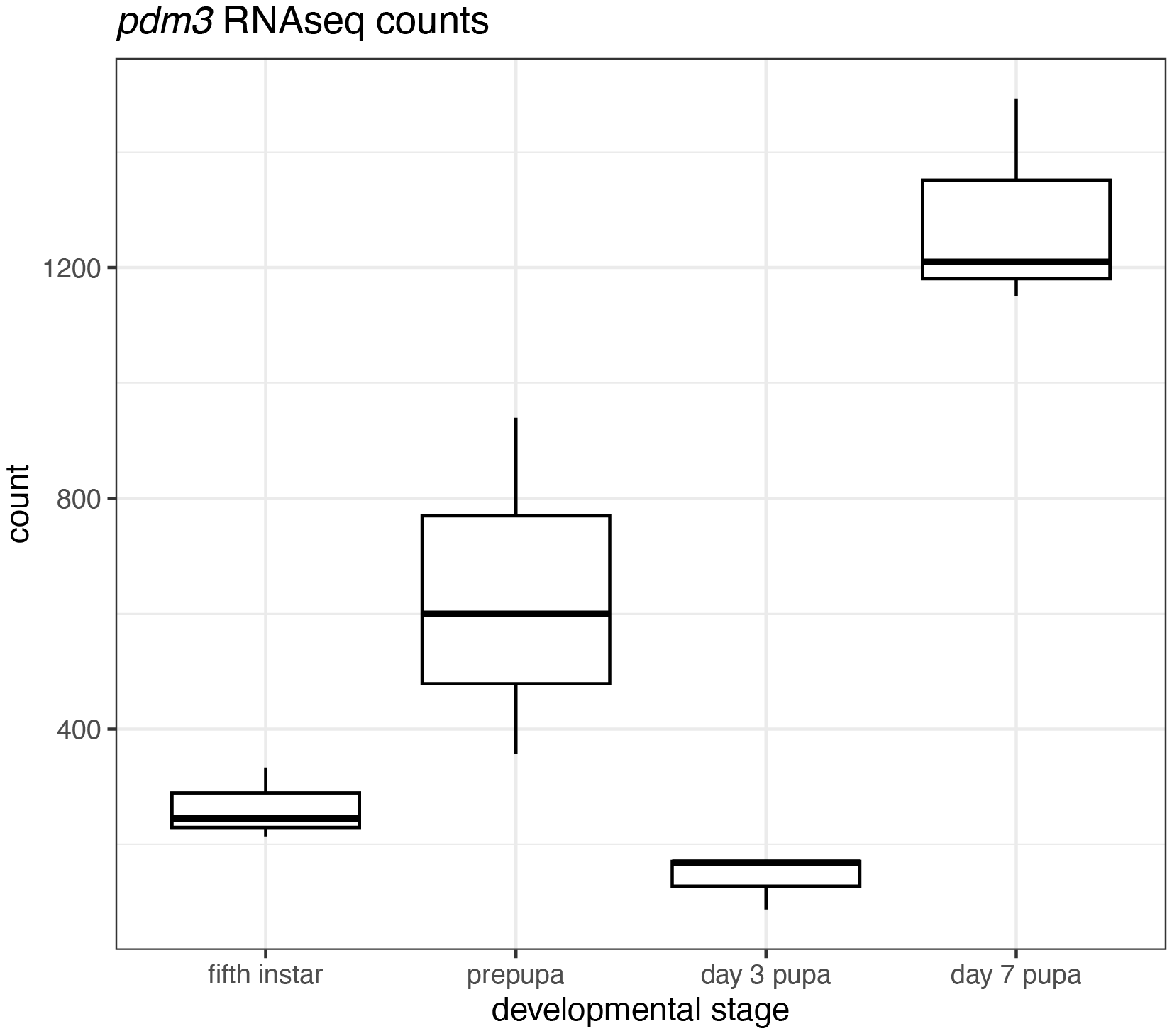

**Fig. S5: Normalized counts of *nubbin* transcripts in the wing across development in *Junonia coenia.***
