## Supplemental File 3 for "*nubbin*, *ventral veinless*, and *pdm3* play diverse roles in butterfly wing pattern development"

**Supplemental File 1: Inference of CRISPR Edits Results**

Indels detected by Synthego’s Inference of CRISPR Edits tool for CRISPR mosaic mutants of each POU domain gene.

***nubbin***

nubbin_02

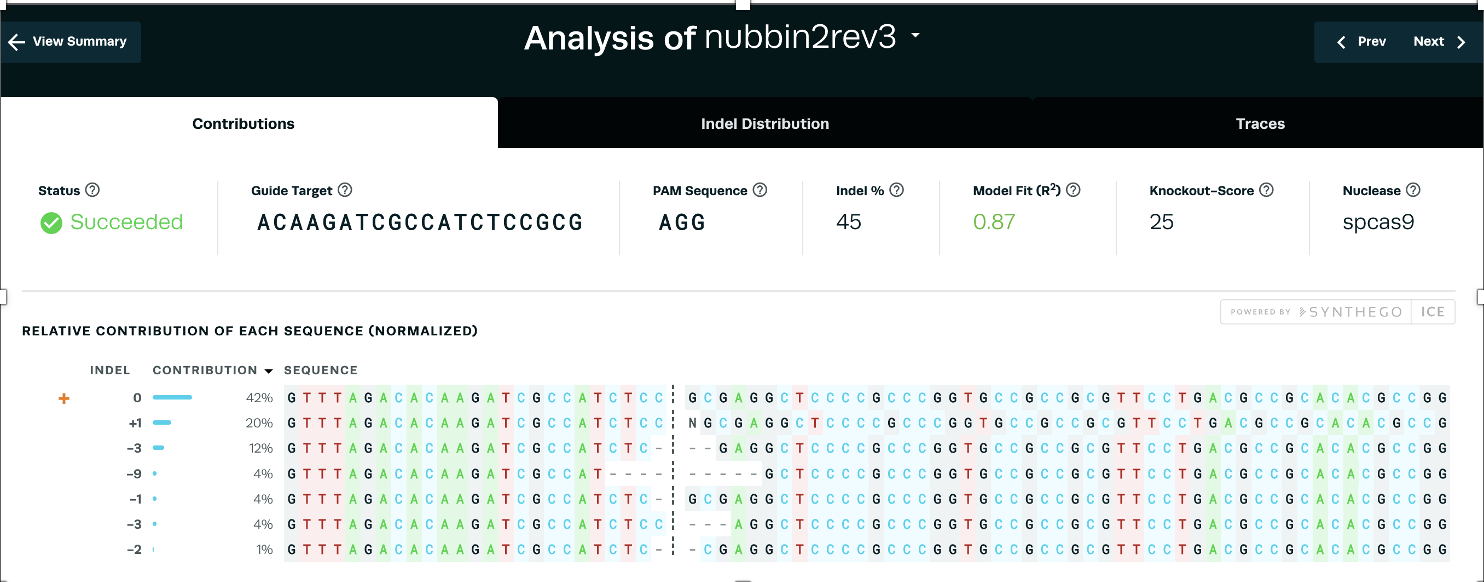

nubbin_04

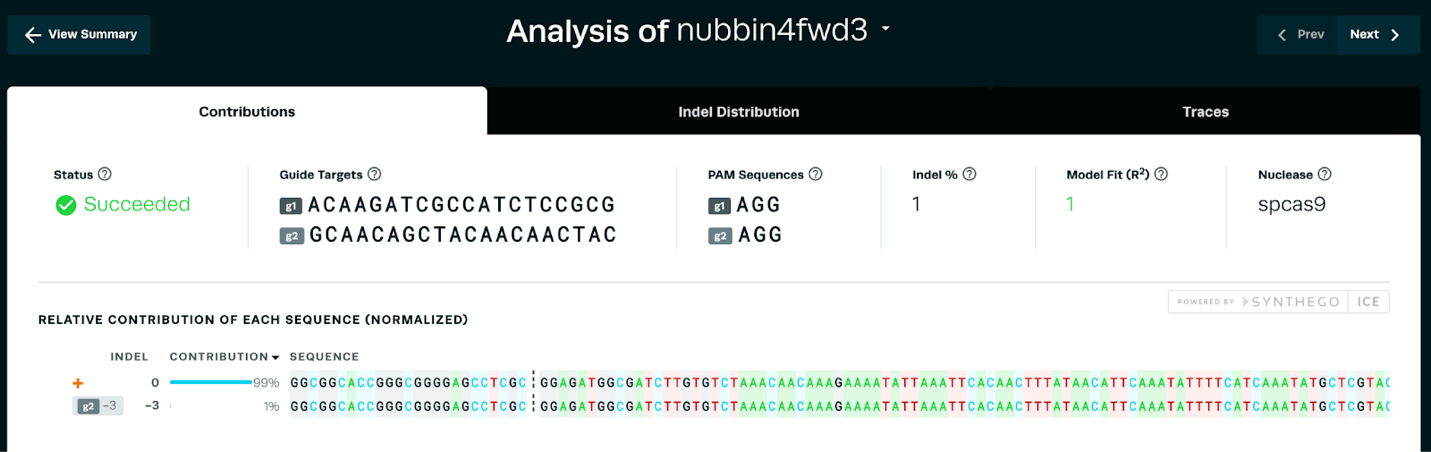

nubbin_67

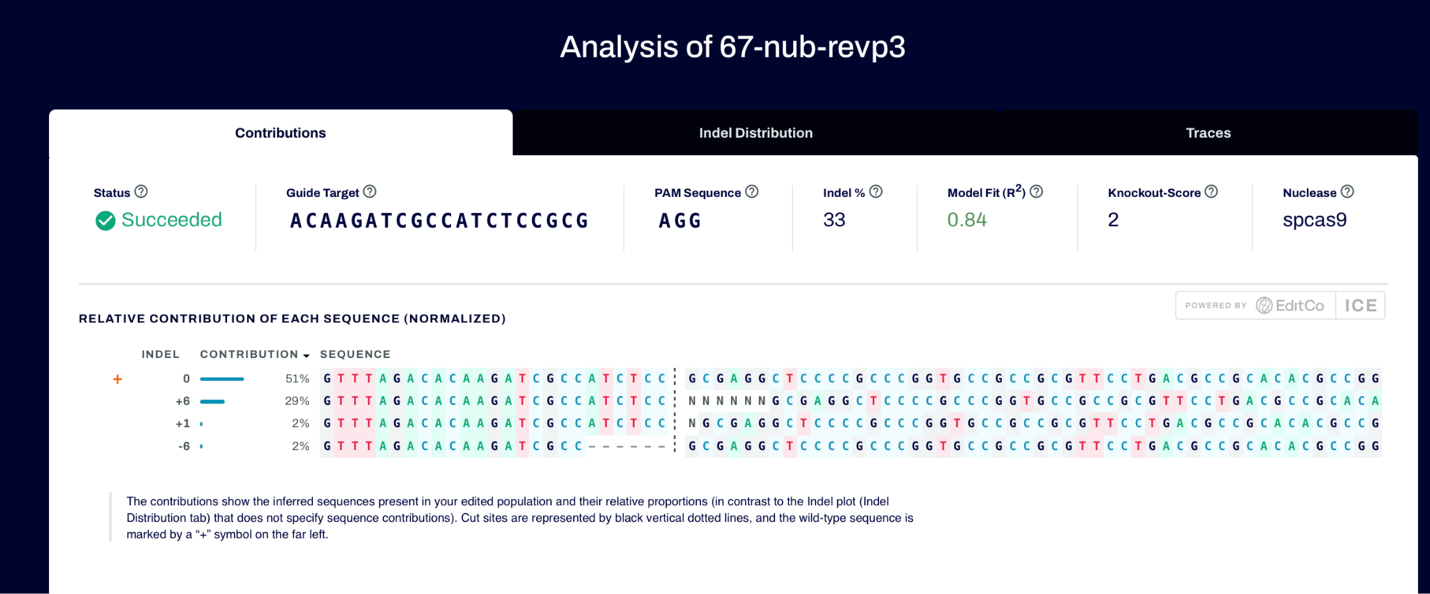

***pdm3***

pdm3_12

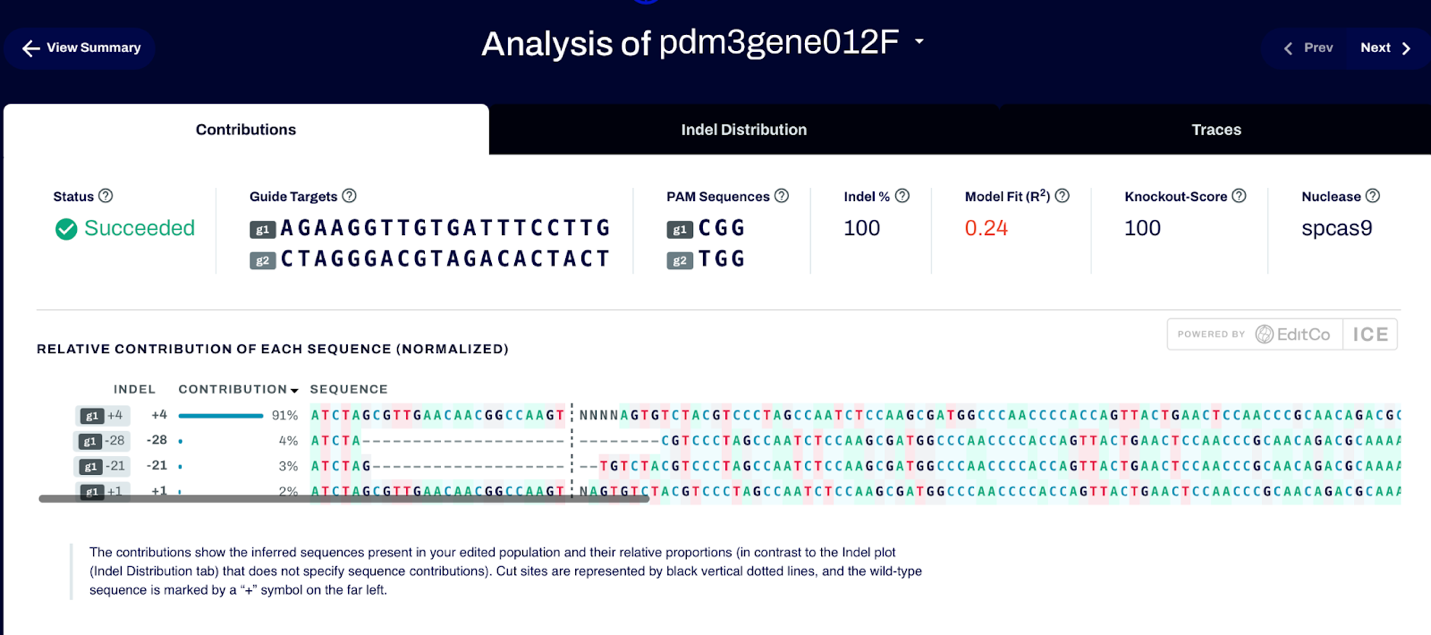

pdm3_45

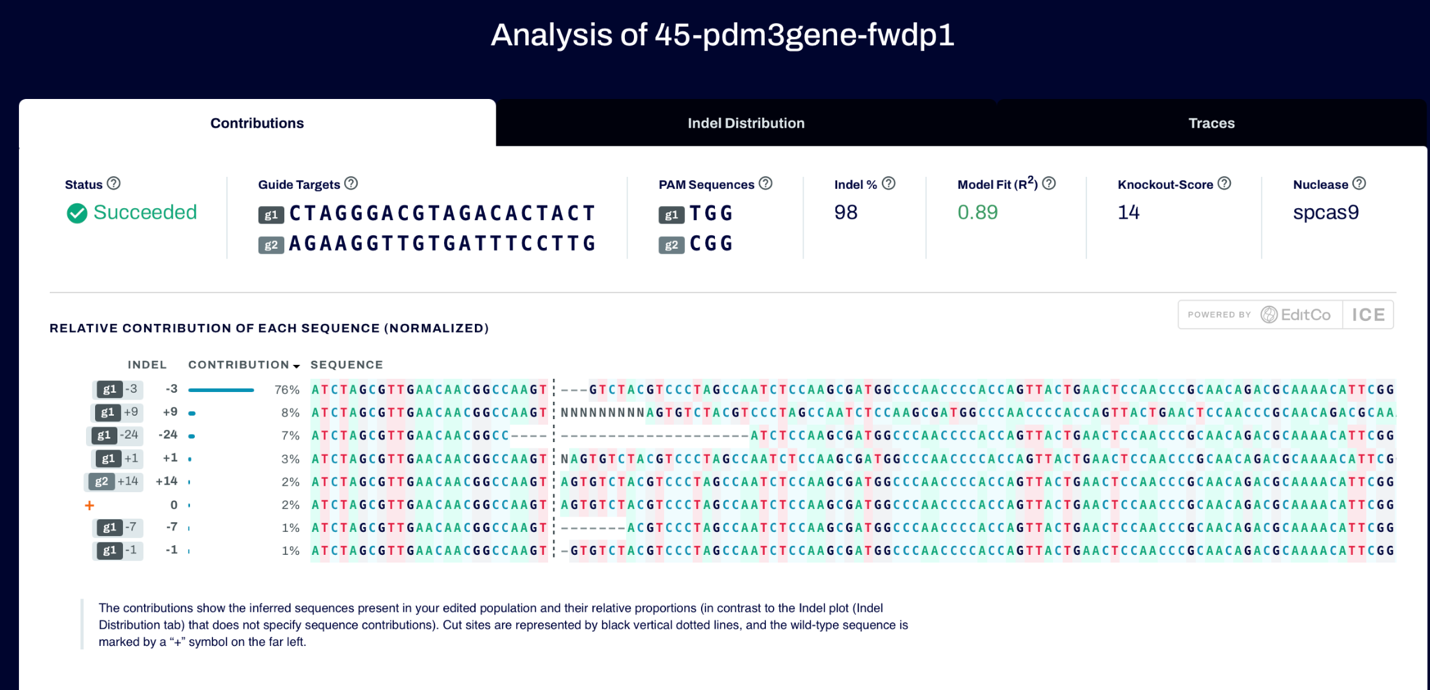

***vvl***

vvl_20
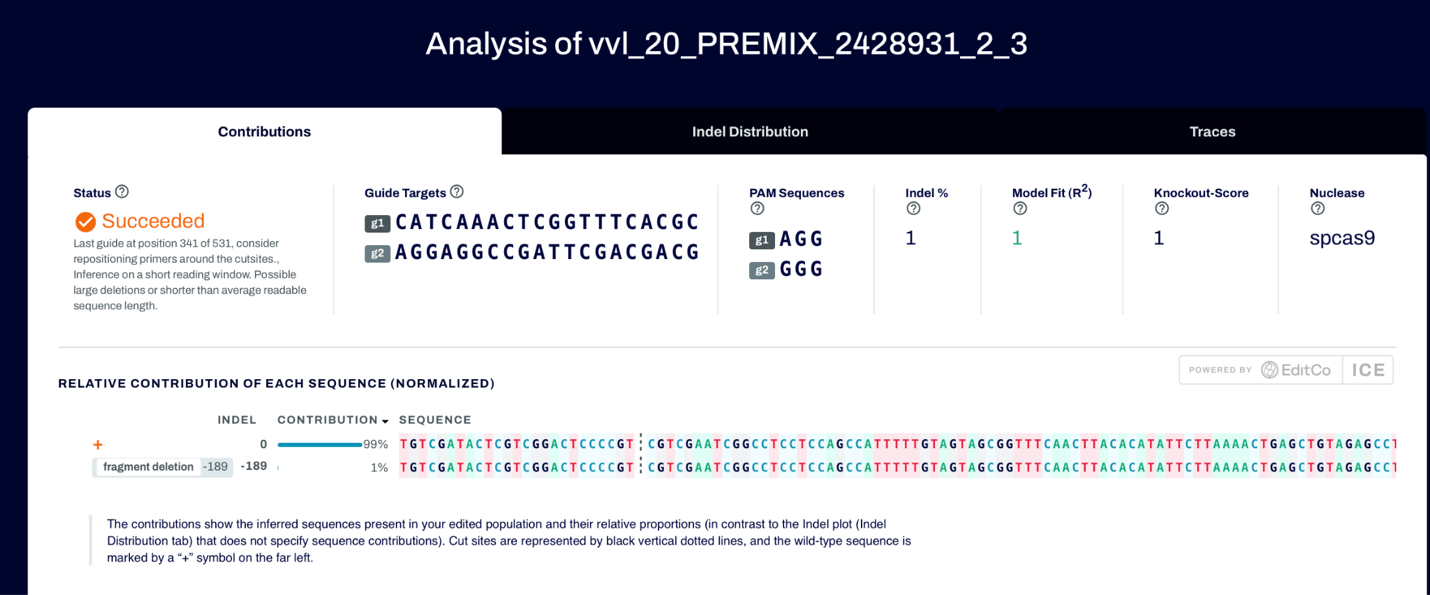
