## Supplemental File 4 for "*nubbin*, *ventral veinless*, and *pdm3* play diverse roles in butterfly wing pattern development"

**Supplemental File 2: Additional HCR in situ hybridization data**

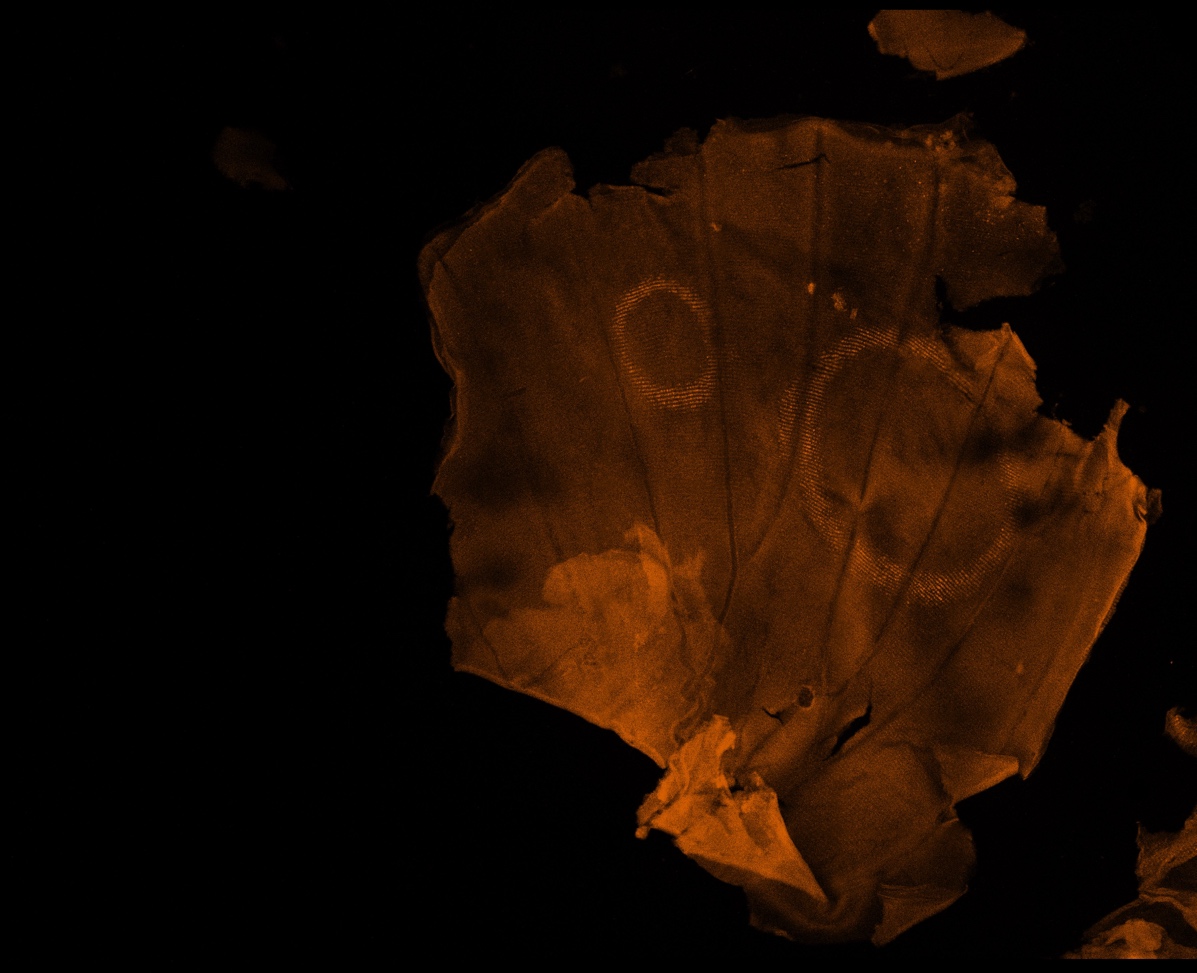

**Fig. S6:** *pdm3* is expressed in the eyespot rings at day 3 of pupal development.

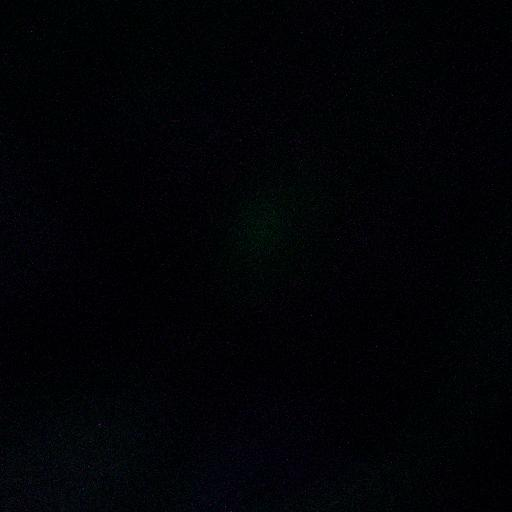

**Movie S1:** Z-stack of *pdm3* (magenta) and *spalt* (green) expression at day 1 of pupal development. Cell nuclei marked with DAPI (blue). (Double click to play video.)

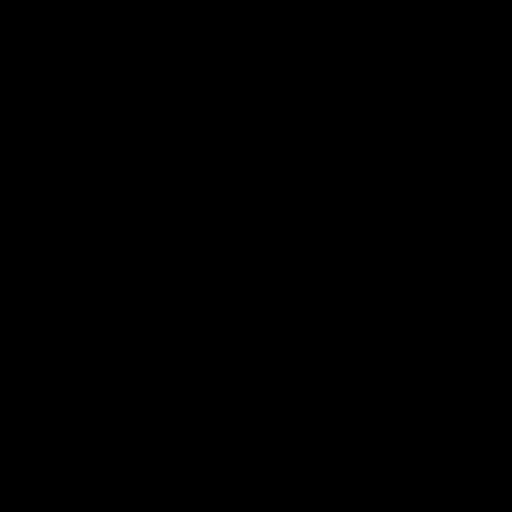

**Movie S2:** Z-stack of *nubbin* (magenta) imaged at 20x in a day 3 pupal wing. (Double click to play video.)
