## Supplementary figures and images for "*nubbin*, *ventral veinless*, and *pdm3* play diverse roles in butterfly wing pattern development"

### Supplemental File 5

***nubbin***

| 01 | 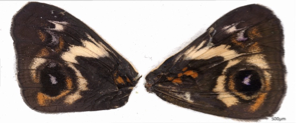  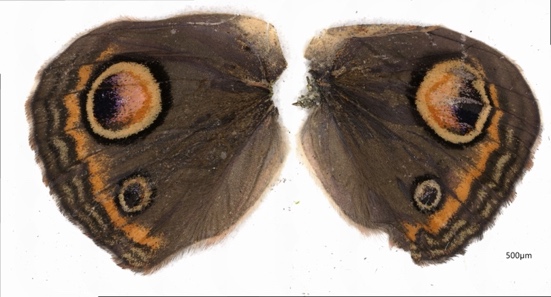 | 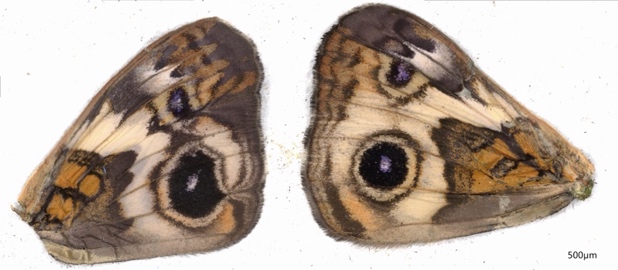  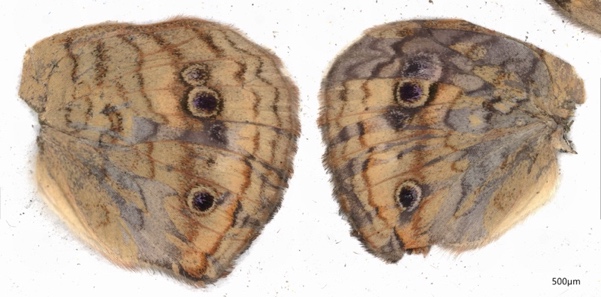 |
| --- | --- | --- |
| 02 | 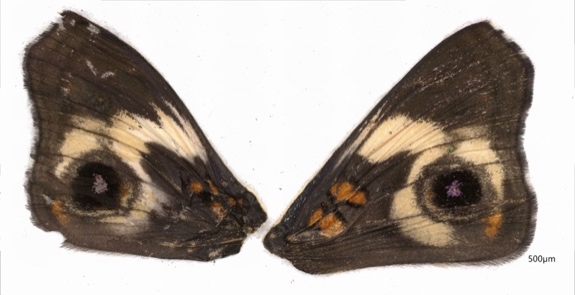  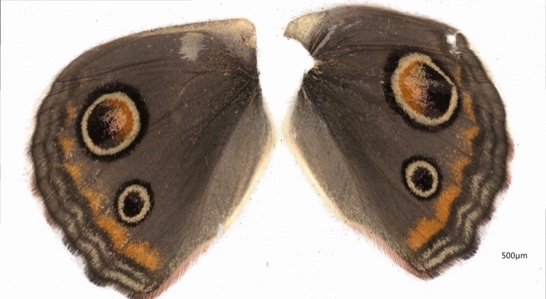 | 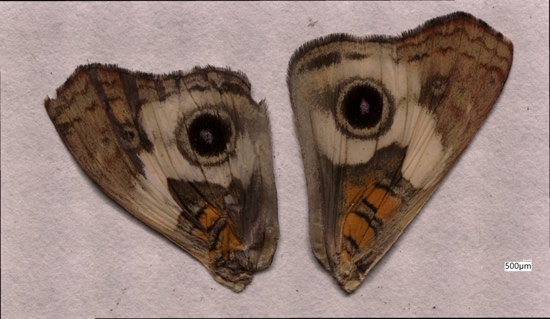  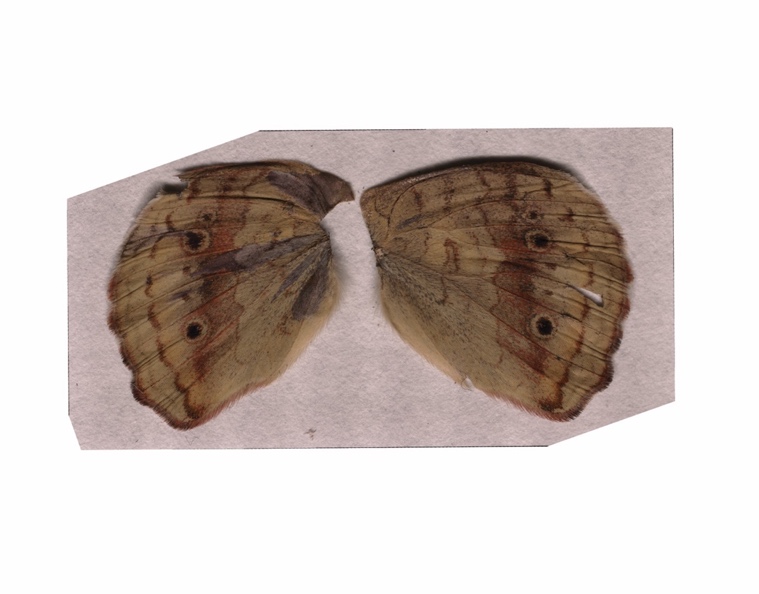 |
| 03 | 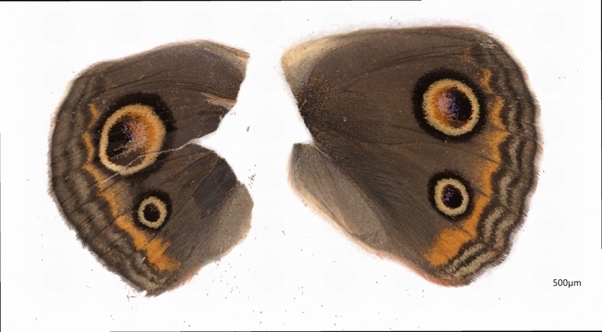 | 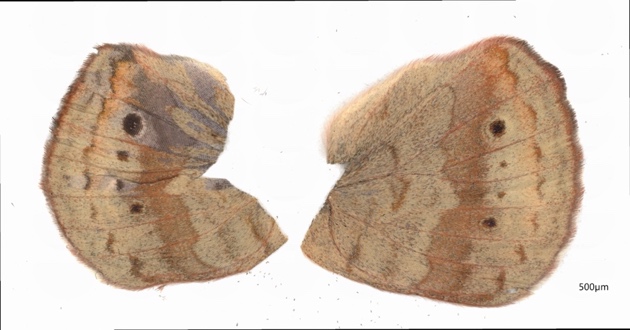 |
| 04 | 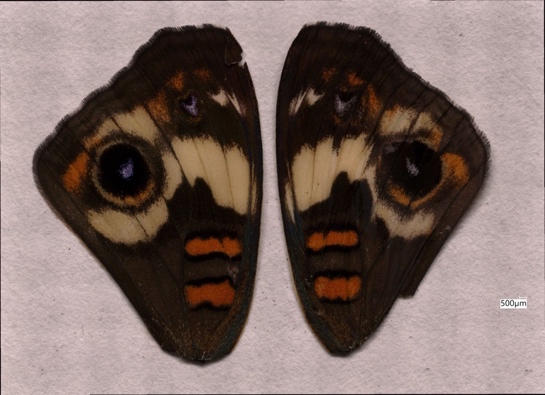 |  |
| 51 | 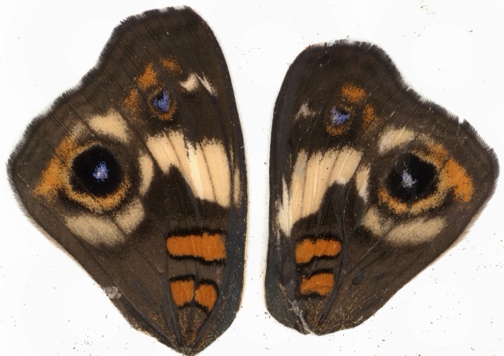  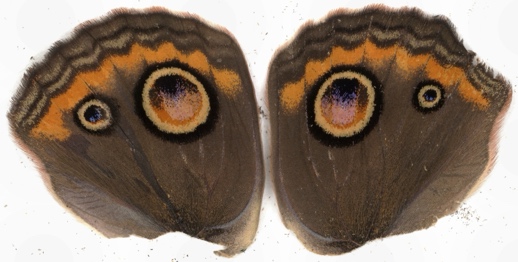 | 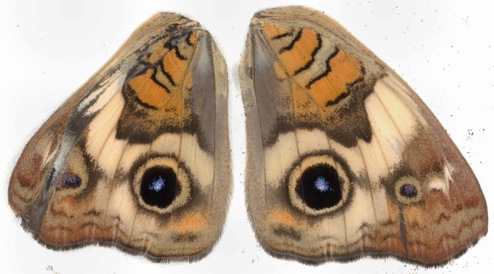  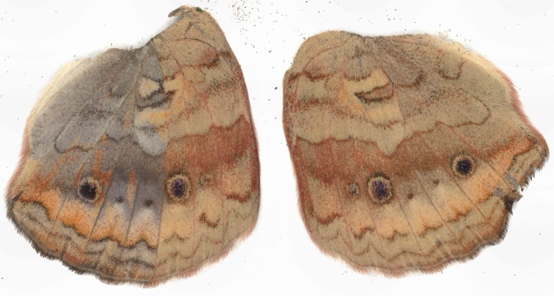 |
| 52 | 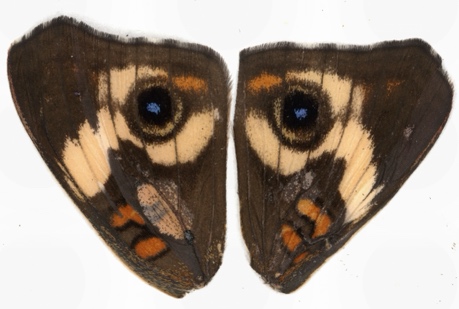  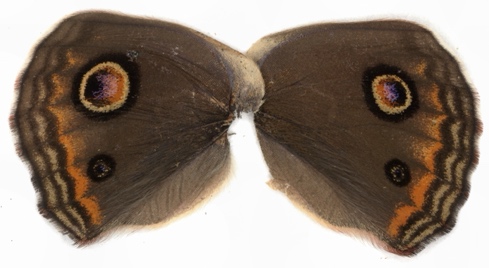 | 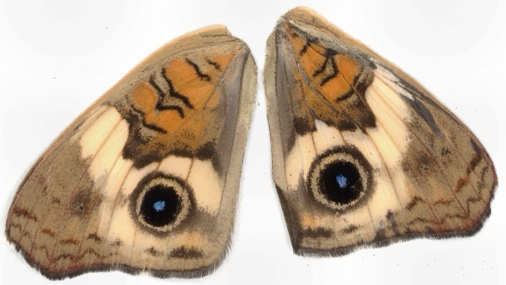  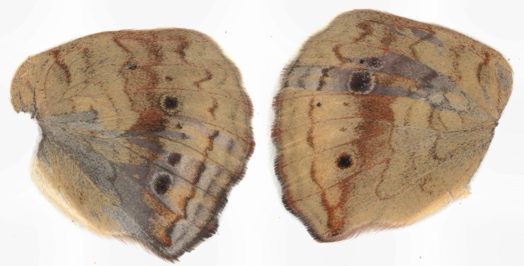 |
| 53 | 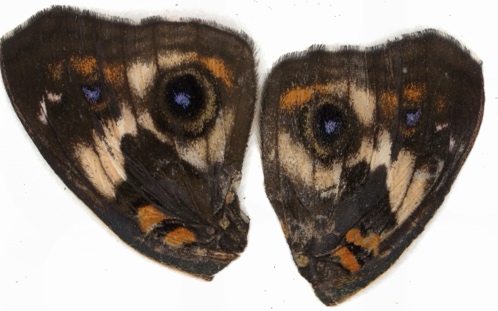  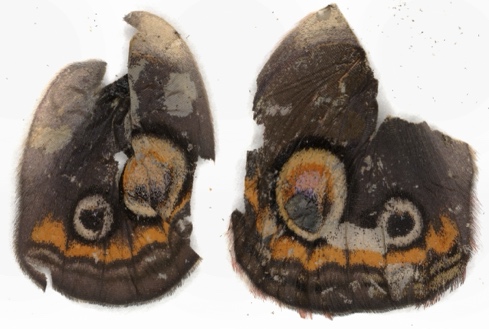 | 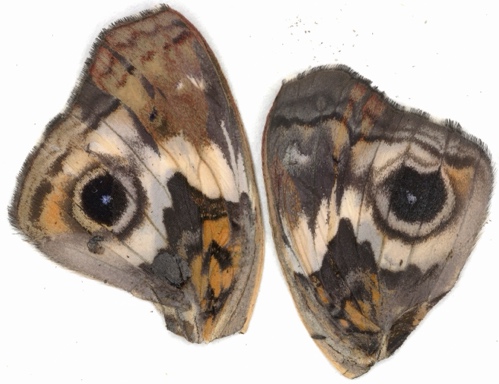  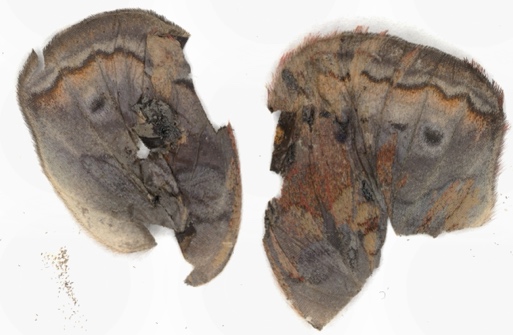 |
| 54 | 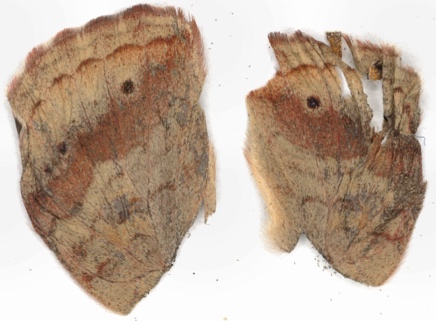 |  |
| 55 | 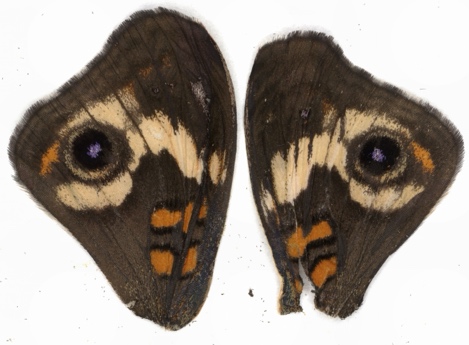  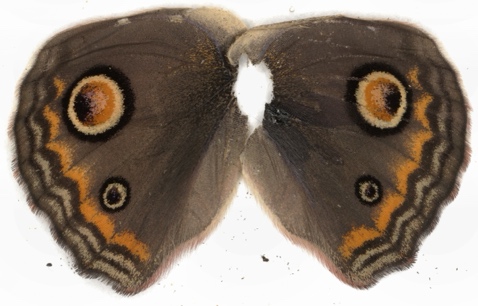 | 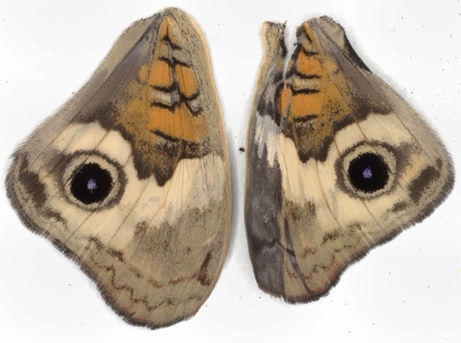  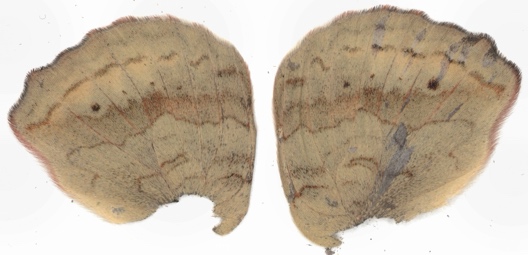 |

| 03  ***vvl*** | 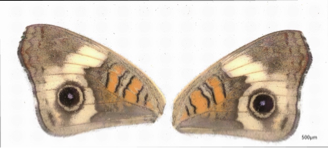 | 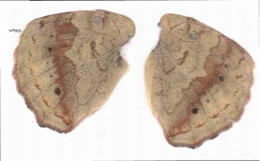 |
| --- | --- | --- |
| 20 | 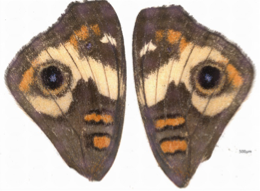 |  |
| 57 | 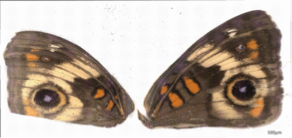 |  |
| 62 | 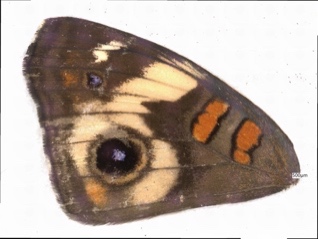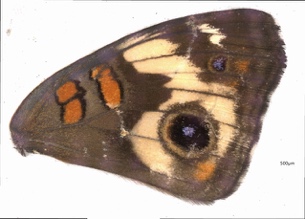 | 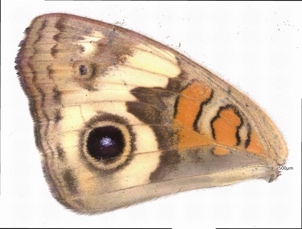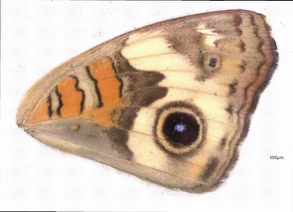 |
| 07 | 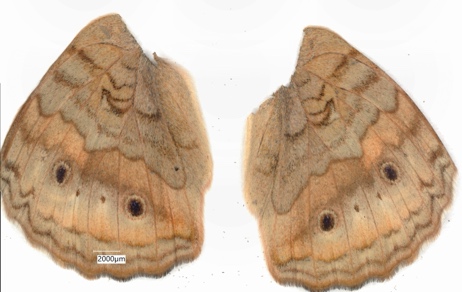 |  |
| 75 | 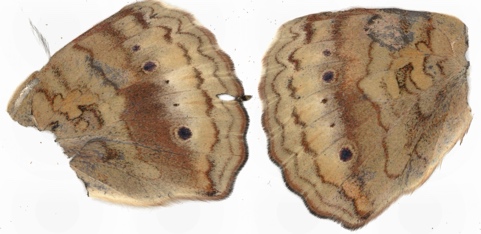 |  |
| 78 | 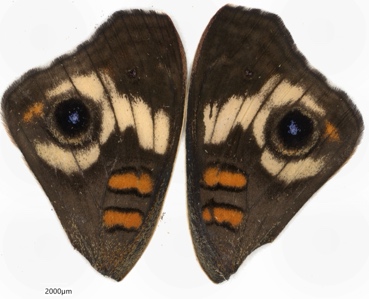 | 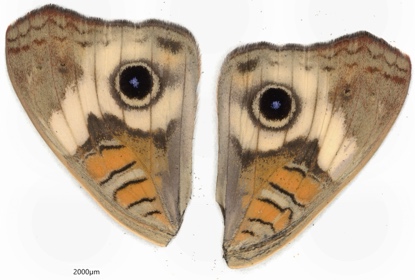 |
| 83 | 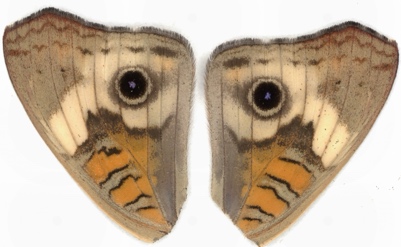 |  |

| 01  ***pdm3*** | 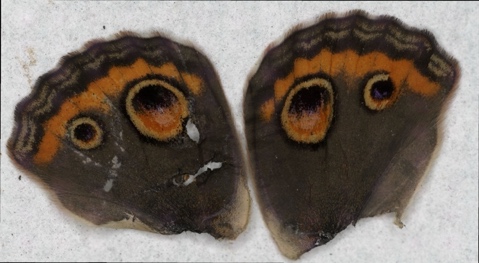 | 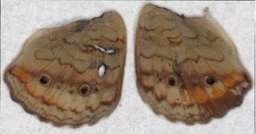 |
| --- | --- | --- |
| 12 | 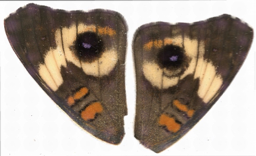  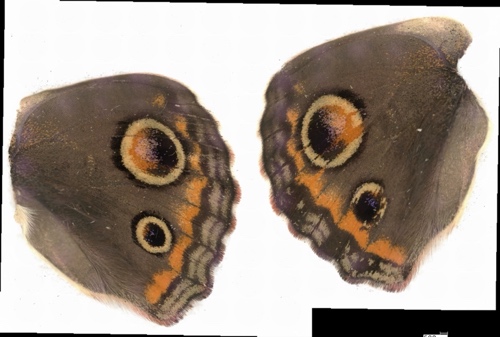 | 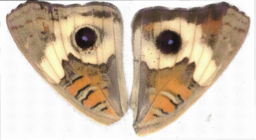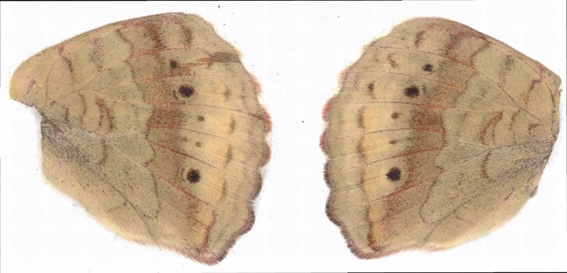 |
| 45 | 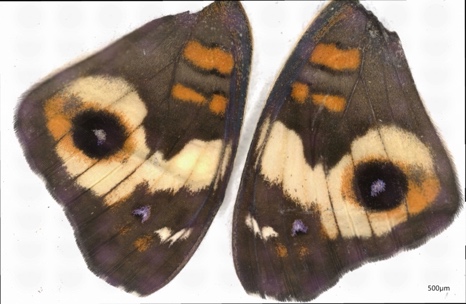  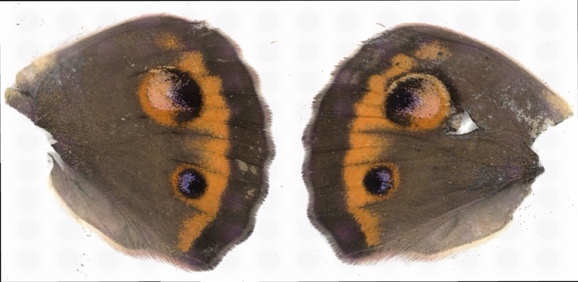 | 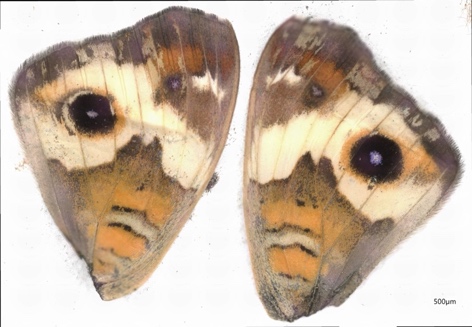  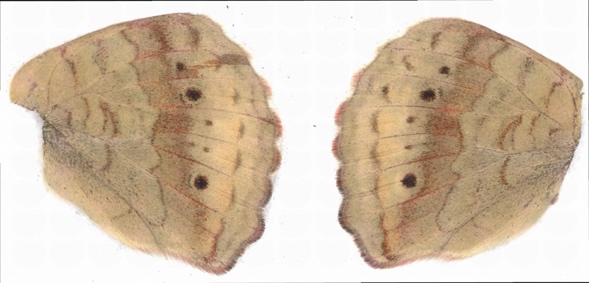 |
| 14 | 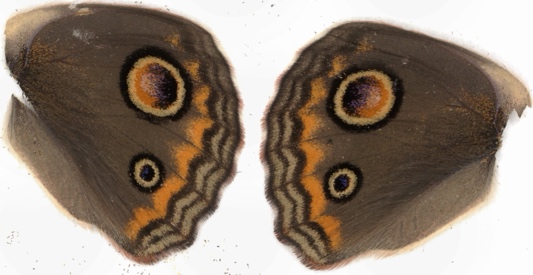 | 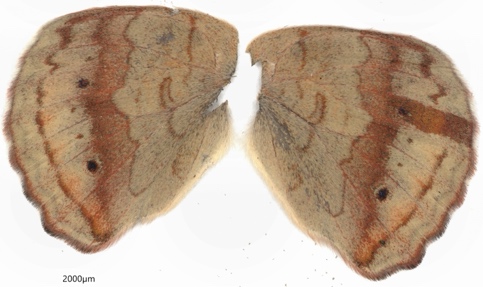 |
| 16 | 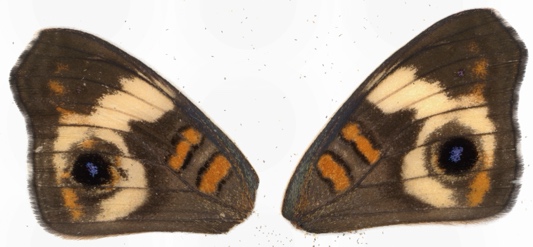  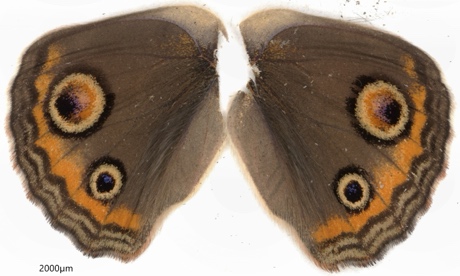 | 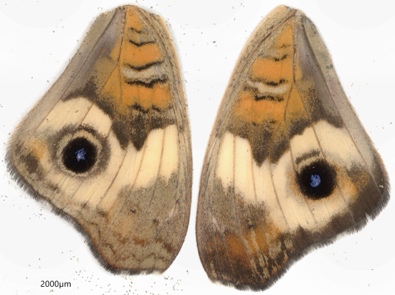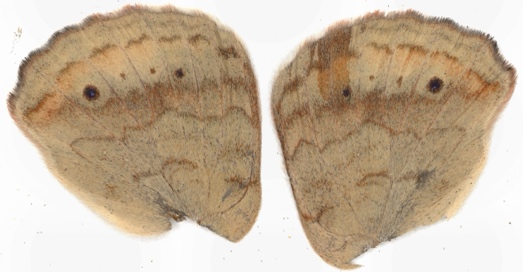 |
| 19 | 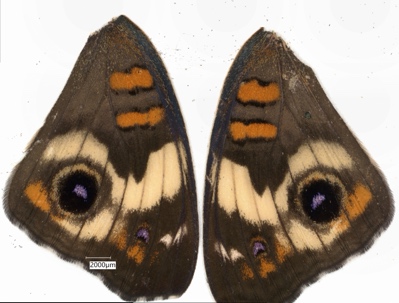  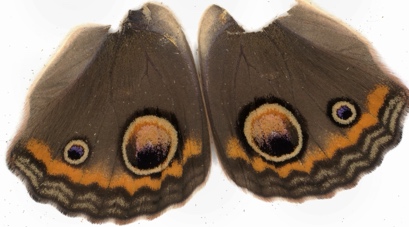 | 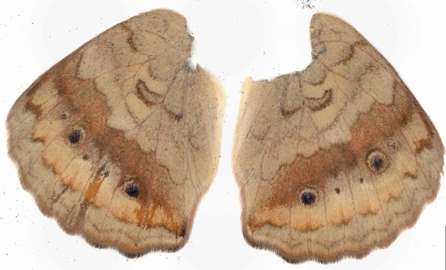 |
| 30 | 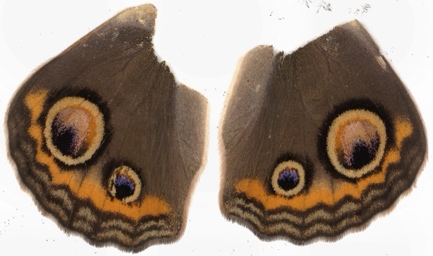 |  |
| 32 | 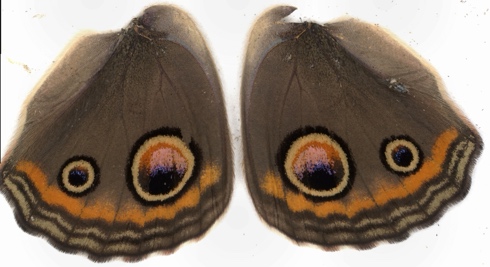 | 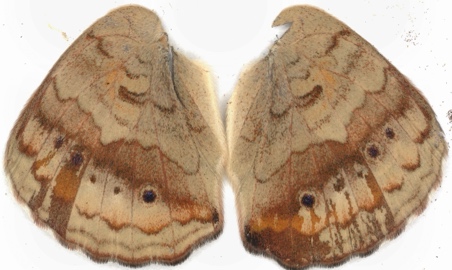 |
| 33 | 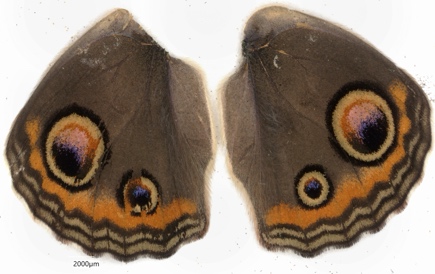 |  |

***pdm3***
